# Accurate Computational Design of 3D Protein Crystals

**DOI:** 10.1101/2022.11.18.517014

**Authors:** Zhe Li, Shunzhi Wang, Una Nattermann, Asim K. Bera, Andrew J. Borst, Matthew J. Bick, Erin C. Yang, William Sheffler, Byeongdu Lee, Soenke Seifert, Hannah Nguyen, Alex Kang, Radhika Dalal, Joshua M. Lubner, Yang Hsia, Hugh Haddox, Alexis Courbet, Quinton Dowling, Marcos Miranda, Andrew Favor, Ali Etemadi, Natasha I. Edman, Wei Yang, Banumathi Sankaran, Babak Negahdari, David Baker

## Abstract

Protein crystallization plays a central role in structural biology^1^, with broad impact^2^ in pharmaceutical formulation^3^, drug delivery^4^, biosensing^5^, and biocatalysis^6,7^. Despite this importance, the process of protein crystallization remains poorly understood and highly empirical^8–10^, with largely unpredictable crystal contacts, lattice packing arrangements, and space group preferences, and the programming of protein crystallization through precisely engineered sidechain-sidechain interactions across multiple protein-protein interfaces is an outstanding challenge. Here we develop a general computational approach to designing three-dimensional (3D) protein crystals with pre-specified lattice architectures at atomic accuracy that hierarchically constrains the overall degree of freedoms (DOFs) of the system. We use the approach to design three pairs of oligomers that can be individually purified, and upon mixing, spontaneously self-assemble into large 3D crystals (>100 μm). Small-angle X-ray scattering and X-ray crystallography show these crystals are nearly identical to the computational design models, with the design target *F*4_1_32 and *I*432 space groups and closely corresponding overall architectures and protein-protein interfaces. The crystal unit cell dimensions can be systematically redesigned while retaining space group symmetry and overall architecture, and the crystals are both extremely porous and highly stable, enabling the robust scaffolding of inorganic nanoparticle arrays. Our approach thus enables the computational design of protein crystals with high accuracy, and since both structure and assembly are encoded in the primary sequence, provides a powerful new platform for biological material engineering.

## Main text

The structures of hundreds of thousands of proteins have been determined by X-ray crystallography^1^. The protein crystals required for this process are typically generated by screening of a wide range of crystallization solution conditions, and are held together by numerous but weak non-covalent interactions (crystal contacts)^10^. Despite the huge amount of effort devoted to protein crystallography, it remains a poorly understood and highly empirical process^8,9^. There has been exciting progress in introducing metal-binding sites^11–13^, crosslinkers^14^, electrostatics^15,16^, DNA^17^, and aromatic interactions^18^ to drive 3D crystal formation, but even in these cases, the atomic structure of the resulting protein material is not directly programmable. A 3D crystal has been computationally designed from short peptides using helix termini as crystal contacts^19^, but despite considerable advances in computational protein design^20^, the general problem of designing sidechain-sidechain interactions across protein interfaces that direct assembly into pre-specified 3D crystal lattices with atomic-level accuracy remains unsolved^21–23^. The problem is challenging because multiple (typically 3 or more) noncovalent protein interaction surfaces with high specificity need to be designed into the monomeric subunits, and each designed interface must have high geometric precision to drive packing into the target crystal lattice–small deviations in interface geometry can add up to large deviations over multiple unit cells and hence disrupt assembly.

## Hierarchical Design Strategy

We set out to develop a divide and conquer hierarchical approach to sequentially design the multiple interfaces required for crystal formation. Crystal space groups in three dimensions are generated from the combination of crystallographic point groups and Bravais lattices^25^. We reasoned that it should be possible to first design interfaces directing assembly of protein monomers into assemblies with point group symmetry, and then, following experimental validation, introduce a third interface constraining the translation of these assemblies to generate the desired Bravais lattice. Here, we focused on space groups containing high symmetry point groups (tetrahedral (T) and octahedral (O)) related by dihedral centers: *P*23, *P*432, *P*4_2_32, *I*432, *F*23, *F*432, and *F*4_1_32^26,27^. In the example shown in Fig. 1a, a diamond lattice of the *F*4_1_32 space group can be constructed by arranging tetrahedral cages formed from two distinct protein monomeric subunits (teal and gray).

**Figure 1.**
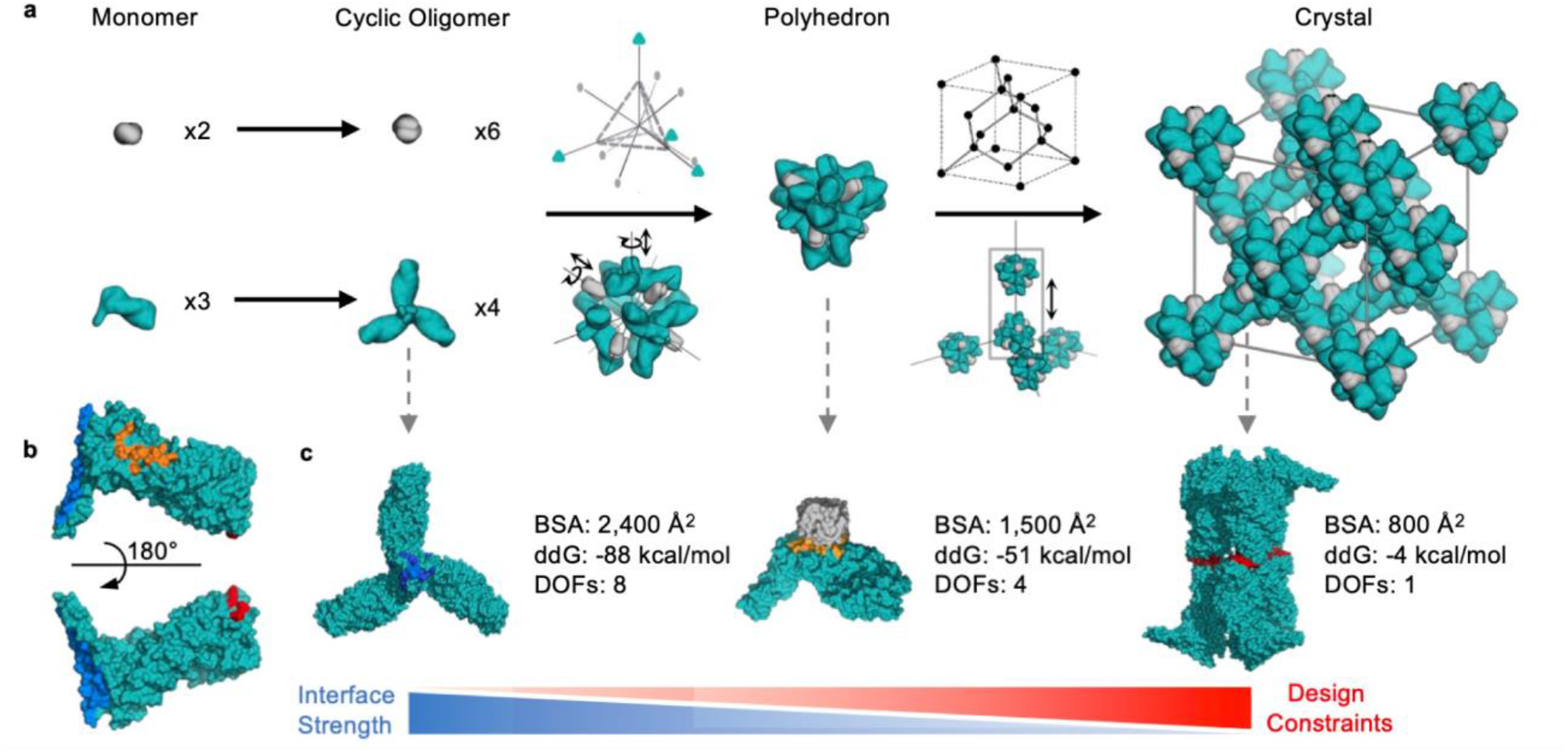
Hierarchical crystal design strategy. **a**, Schematic illustration of the three step design hierarchy for a diamond lattice (*F*4_1_32 space group) formed from a tetrahedral polyhedron built from a C2 dimer (grey) and a C3 trimer (cyan). Monomers (first column) are docked into cyclic dimers and trimers (second column) which are docked into a two-component cage (third column) which is then arrayed in a 3D lattice (fourth column). **b**, Interfaces driving crystal assembly. For the designed crystal example in a, the three designed interfaces on the trimer component are shown that drive assembly of the cyclic oligomer (blue), the tetrahedron (orange), and the crystal lattice (red). **c**, Interfaces mapped to the monomer in b) are shown between interacting partners for the trimer (left), tetrahedral cage (middle) and crystal (right). To maximize assembly cooperativity, the interface size (buried surface area, BSA) and affinity (Rosetta calculated ddG^24^) decrease through the design hierarchy; the number of system degrees of freedom (DOFs) available for sampling at each step (see methods) decreases in parallel making the design challenge more difficult.

The successful implementation of this design strategy requires three orthogonal interfaces on the same protein component (Fig. 1b) with decreasing interface strength at each assembly step to maximize assembly cooperativity, and with high geometric precision to satisfy the overall crystal lattice symmetry constraints (Fig. 1c). Hierarchical design of the first two interfaces has been achieved previously^28–31^, but the design of the third interface (Fig. 1c, right panel shows interface formed between two tetrahedra interacting through two trimers adopting a dihedral D3 arrangement) is an unsolved challenge as only a single translational DOF is available for sampling, making it difficult to generate favorable shape-complementary backbone arrangements for sequence design (the targeted *F*4_1_32 space group requires that the C2 axes of the D3 dihedral and the crystal be coincident; Extended Data Fig. 1a)). To address this issue, during the point group symmetry construction stage, we implement geometric filters to ensure sufficient secondary structure interaction between the trimer building blocks along the direction of crystal propagation to facilitate the design of specific sidechain-sidechain interactions across the interface (Extended Data Fig. 1b). The Bravais lattice generating interface must specifically encode the desired crystal lattice, and be orthogonal to the cage and oligomer interfaces to not interfere with cage assembly or promote off-target assemblies. In addition to proper backbone docking geometry, the crystal contact interface must be in a narrow window of interaction strength (Extended Data Fig. 1c): if too strong, there will be little disassociation between interacting cages and hence limited self-correction of assembly errors, resulting in kinetic traps; if too weak, there will be insufficient driving force for assembly of the target crystal structure.

To implement this crystal design strategy, we designed new protein polyhedra favorable for assembly into crystals (Extended Data Fig. 2, e-l, Extended Data Fig. 3, Extended Data Fig. 4) by using RPXDock^32^ to dock a wide range of designed oligomers with C2, C3 or C4 symmetry into T33, T32, O43, O42 and O32 assemblies (the two indices indicate the symmetries of the constituent oligomers; O43 indicates an octahedron generated from a designed cyclic tetramer and cyclic trimer) and designing sequences directing assembly using Rosetta^24^. Following experimental validation, the polyhedra were placed into the corresponding Wyckoff sites in the target crystal lattice, the translational spacing sampled finely around 3 Å of close contact (methods, Extended Data Fig. 5), and the sequence at the interface designed using Rosetta to direct crystal assembly aiming for specific but lower affinity interactions than within the polyhedra to increase assembly cooperativity.

## Computational Design and Characterization of *F*4_1_32-1, *F*4_1_32-2 and *I*432-1 Crystals

To independently evaluate the last step in our hierarchical approach, we first sought to assemble previously designed tetrahedral cages^29,33,34^into *F*4_1_32 lattices through a newly designed D3 symmetry cage-cage interface. In a first round of design calculations, all high scoring designs were found to employ the same building block, T33-15^29^, a 24-subunit tetrahedral cage composed of two distinct cyclic trimeric building blocks C3-A and C3-B, with C3-B forming the extended crystal interface. The T33-15 cage can be assembled *in vitro* from separately purified components^29^, which enabled facile screening of crystal designs by mixing C3-B containing the crystal interface with the unmodified C3-A *in vitro* at a 1:1 molar ratio in hanging drops (Method). For the design T33-15-D3-4, we observed octahedral-shaped crystals up to 200 μm in diameter (F4_1_32-1-0) over a week (Extended Data Fig. 6a). With a Thr to Arg substitution at the periphery of the crystal interface, crystals formed within 12 hours of mixing without additional precipitant; we refer to this crystal as *F*4_1_32-1 (Fig. 2b inset, Extended Data Fig. 6b). To examine the structure of the constituent cage without the complications of rapid crystal assembly, we characterized T33-15-D3-4 prior to crystallization by size exclusion chromatography (SEC) (Extended Data Fig. 2a), negative stain electron microscopy (nsEM) (Extended Data Fig. 2, b-c), and small-angle X-ray scattering (SAXS) (Extended Data Fig. 2d) which were consistent with the designed cage structure. To study the crystal packing and molecular details of the *F*4_1_32-1 crystal interface, we characterized the *F*4_1_32-1 crystals by cryoEM (Fig. 2b, Extended Data Fig. 6c), SAXS (Fig. 2c) and X-ray crystallography (Fig. 2d). SAXS data collected at room temperature indicated *F*4_1_32 symmetry with a unit cell diameter of 295 Å, in close agreement with the designed lattice model (297 Å), and the experimental scattering intensity profile closely matched that computed from the designed crystal lattice (Fig. 2c). A 2.80 Å resolution X-ray structure of the crystal showed that both the overall designed *F*4_1_32 lattice and the individual subunit-subunit interfaces were closely recapitulated (Fig. 2d). The all-atom root-mean-square deviation (RMSD) between the X-ray structure and computational design model over the six monomers that form the dihedral tetrahedron-tetrahedron interface was 1.19 Å (Extended Data Fig. 7b), and over two full contacting cages, 1.39 Å (Extended Data Fig. 7c). The crystal structure of the original T33-15-D3-4 design (prior to the Thr55Arg substitution, Supplementary Fig. 1) has a slightly higher RMSD (1.81 Å) to the design model, likely due to the formation of an unintended hydrogen bond by the Thr (Supplementary Fig. 2).

**Figure 2.**
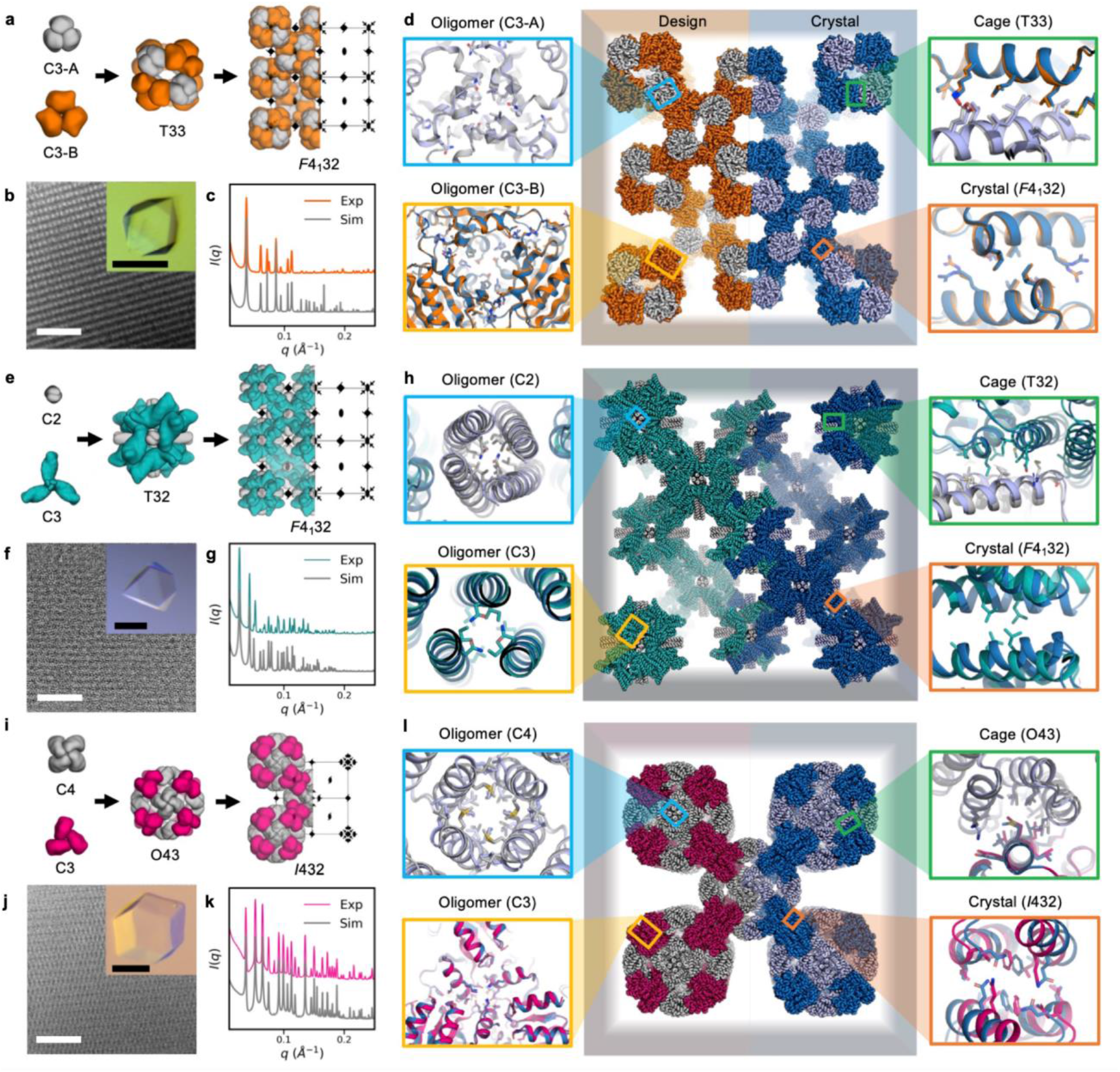
Computational design and experimental characterization of *F*4_1_32-1 and *I*432-1 crystals. **a-d**, *F*4_1_32-1, **e-h**, *F*4_1_32-2, **i-l**, I432-1. **a**,**e**,**i**, Construction of crystals from cyclic oligomers. In the second step, symmetry elements of the cage are superimposed with corresponding symmetry elements of the unit cell. **b**,**f**,**j**, CryoEM images of crystals (scale bar, 100 nm) and optical micrograph of single crystals (inset, scale bar, 100 μm). **c**,**g**,**k**, SAXS spectra of microcrystals (orang, teal and hot pink) compared to theoretical spectra computed from the design model (gray); there is a close agreement in all three cases. **d**,**h**,**l**, Middle panel, computational design model (left, orange, teal, hot pink and gray) spliced to experimentally determined crystal structure (right, sky blue and light blue). The four flanking panels show zoom-ins on the four designed interfaces, with the design model superimposed on the crystal structure. Left top and bottom: the two cyclic oligomer interfaces; Top right; interface between cyclic oligomers that generates polyhedral cage; Bottom right, interface between polyhedra that generates the crystal. All-atom RMSDs between design model and experimentally determined structure for each interface for crystal design *F*4_1_32-1 are C3-A: 0.42 Å; C3-B: 0.93Å; T33: 0.73 Å; *F*4_1_32: 1.19 Å; for crystal design *F*4_1_32-2, C2: 1.73 Å; C3: 1.98 Å; T32: 1.86 Å; *F*4_1_32: 3.00 Å, and for crystal design I432-1, C4: 1.10 Å; C3: 1.00 Å; O43: 1.21 Å; *I*432: 1.34Å.

While the *F*4_1_32-1 crystal design demonstrated the capability of our computational protocol in programming assembly to pre-specified crystal lattices with angstrom level resolution, it also highlighted some remaining challenges. First, the fine balance between interface affinity and specificity required to drive the assembly process while avoiding off-target interactions (Extended Data Fig. 1c); many designs were insoluble or formed off-target assemblies likely due to overly extensive designed crystal interfaces (Supplementary Fig. 3). Second, the previously characterized polyhedral protein assemblies only allowed limited exploration of the crystal design space because the constituent oligomeric building blocks in most cases did not present designable secondary structure elements capable of forming the new crystal interface (Extended Data Fig. 1b). Hence, we designed a new set of polyhedral cages specifically for crystal design and used them as building blocks for crystals (Extended Data Fig. 2, e-l, Extended Data Fig. 3, Extended Data Fig. 4).

**Figure 3.**
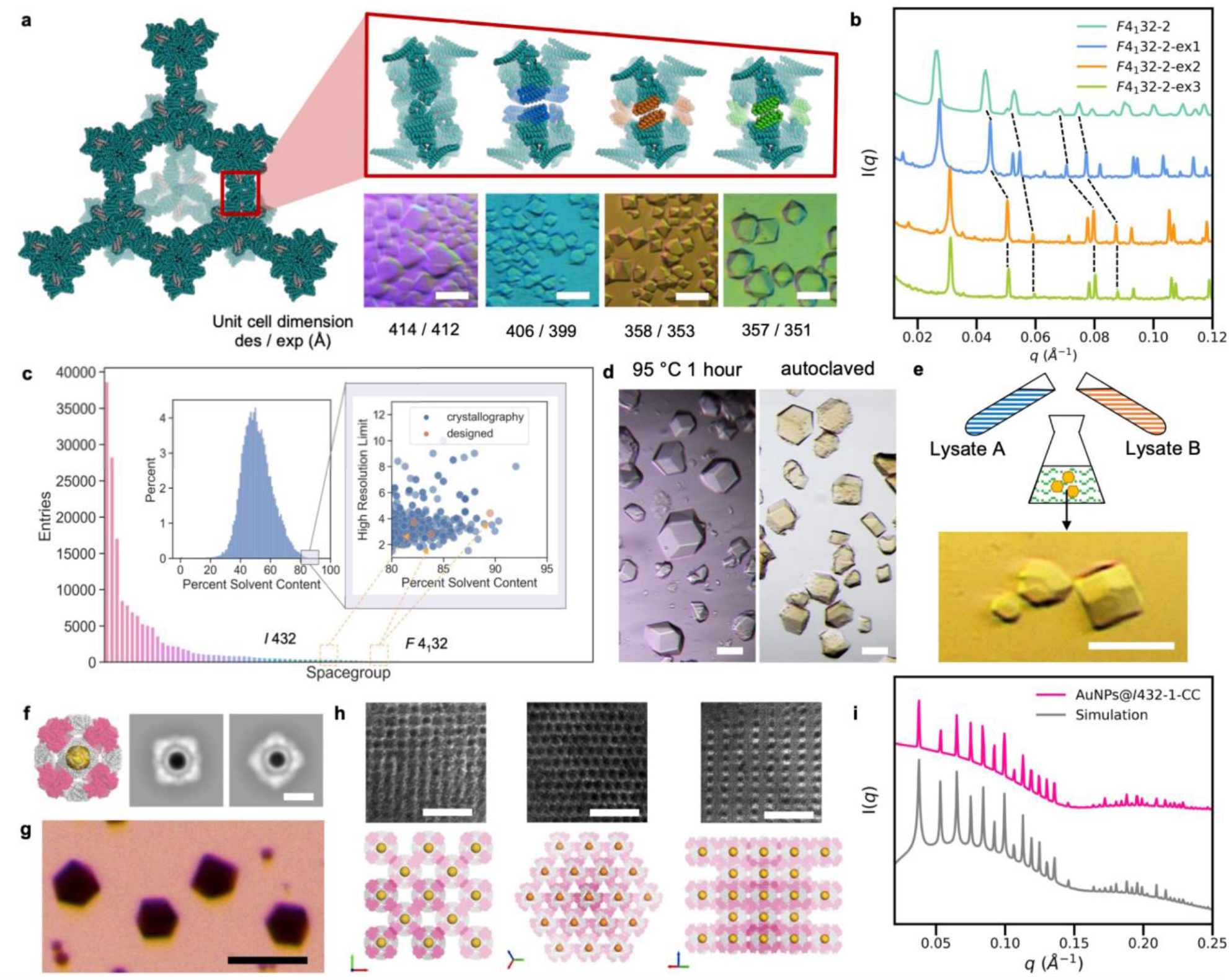
Engineering crystal properties. **a**,**b**, Tuning the unit cell dimensions of the *F*4_1_32-2 crystals by design. **a**, Design models and optical micrographs. Scale bar, 50 μm. **b**, SAXS profile. Peaks of the same index are connected by dash lines. **c**, Space group distribution over all crystals in the PDB. Inset: distribution of crystal solvent content, with zoom-in on high solvent content region versus crystal resolution. Designed crystals in this paper are highlighted in orange. **d**, *I*432-1-CC crystals incubated at 95 °C for 1 hour (left panel) and autoclaved at 121 °C, 13 psi for 40 mins (right panel). Scale bar, 100 μm. **e**, *I*432-1-CC crystals formed overnight by mixing of *E. coli* lysates. Scale bar, 50 μm. **f-i**, Scaffolding 3D gold nanoparticle (AuNP) arrays by *I*432-1-CC crystals. **f**, Model of O43-2 cage with AuNP encapsulation (left panel) and representative 2D class averages of nsEM images (right panel). Scale bar, 10 nm. **g**, Optical micrograph of crystals with AuNP encapsulation. Scale bar, 50 μm. **h**, NsEM micrographs (top panel) and 2×2×2 unit cell crystal model (bottom panel) of different crystals facets: <100> (left), <111> (middle) and <110> (right). Scale bar, 50 nm. **i**, Experimental SAXS profiles of protein crystal scaffolded AuNP arrays (pink) and simulated pattern (grey) of superlattice formed by 5.6 nm diameter AuNPs with a lattice parameter of 236.3 Å. Owing to a significantly lower scattering cross-section of proteins, as compared to that of the AuNPs, no distinguishable scattering intensity was observed from the protein host lattice.

We set out to design a new diamond lattice using the custom designed polyhedral building blocks (Fig. 2e). We first designed a T32 cage from a pH-responsive trimer^35^ rigidly fused to designed helical repeat proteins (DHRs)^36^ (HFuse_pH192_0046, manuscript in preparation) and a helical bundle dimer (2L4HC2_23)^30^ (Supplementary Fig. 4a). We designed this cage, T32-15, such that when docked into a diamond lattice it forms dihedral D3 crystal contact interfaces at the facets defined by the centers of three neighboring trimers along the other C3 symmetry axis of the tetrahedron (Fig. 1c, right panel), as opposed to the vertices defined by a single trimer as in the *F*4_1_32-1 design (Extended Data Fig. 1b, top panel). We confirmed cage formation by immobilized metal affinity chromatography (IMAC) pull-down followed by SDS-PAGE analysis (Supplementary Fig. 4b), SEC (Extended Data Fig. 2e), nsEM (Extended Data Fig. 2f, Supplementary Fig. 4c), and SAXS (Extended Data Fig. 2h), and determined the structure by Cryogenic electron microscopy (cryoEM) with a global 3.34 Å resolution (Extended Data Fig. 2g, Extended Data Fig. 8). The cryoEM structure closely matched the design model over both designed components, with some deviation in helix orientation of the C3 component at the periphery of the cage (Extended Data Fig. 2g). Genes encoding six designed lattices were obtained, and after mixing the independently purified components, design T32-15-D3-6 formed octahedral-shaped crystals (*F*4_1_32-2-6H, Extended Data Fig. 6d). We found that shortening the helices of the C2 component increased its solubility and the yield of cage assembly following mixing (Supplementary Data Fig. 5). This trimmed cage, T32-15-D3-6-6H, formed ~100 μm sized octahedral-shaped crystals (*F*4_1_32-2) in 3 to 4 days (Fig. 2f inset, Extended Data Fig. 5e). Crystals were characterized by cryoEM (Fig. 2f, Extended Data Fig. 6f), SAXS (Fig. 2g) and X-ray crystallography (Fig. 2h). Indexing of the experimental SAXS spectrum of *F*4_1_32-2 microcrystals indicated the desired *F*4_1_32 space group with a unit cell edge length of 412 Å, in close agreement with the designed lattice model (417 Å, Fig. 2g). We solved the crystal structure of the designed lattice to 4.40 Å using the cryoEM model of the cage (Extended Data Fig. 8) for molecular replacement (Fig. 2h). The experimentally determined crystal lattice is very similar to the computationally designed model both in overall structure and at the individual crystal lattice interfaces, with all-atom RMSDs of 3.00 Å over the full dihedral (Extended Data Fig. 7e) and 3.77 Å over two complete interacting cages (Extended Data Fig. 7f). The lower resolution of *F*4_1_32-2 likely reflects intrinsic flexibility at the crystal contacts; the T32-15 cage cryoEM map had a lower local resolution (4~5 Å) at the periphery of the helical fusion region (Extended Data Fig. 8). The *F*4_1_32-2 crystal is, to our knowledge, the first example of a macroscopic, three-dimensional crystalline material computationally designed from bottom-up with all *de novo* proteins and with high accuracy.

To investigate whether our crystal design approach could be applied to a broader range of space groups, we sought to design *I*432 crystals using designed octahedral cages to occupy the constituent Wyckoff sites (Fig. 2i). We designed cage O43-2 from a hyperthermophilic TIM barrel trimer (PDB ID:1WA3) and a *de novo* designed helical repeat protein tetramer (tpr1C4_2) ^28^ (Supplementary Fig. 4d); the 1WA3 trimer satisfies the design rule for presenting accessible secondary structures for the design of crystal contacts (Extended Data Fig. 1b). The two components could be readily expressed and purified separately, and they assembled *in vitro* to form cages at high yield as evidenced by IMAC pull-down followed by SDS-PAGE analysis (Supplementary Fig. 4e), SEC (Extended Data Fig. 2i), nsEM (Extended Data Fig. 2, j-k, Supplementary Fig. 4f) and SAXS (Extended Data Fig. 2l). One of the six ordered designs, O43-2-D3-6, formed rhombic dodecahedral shaped crystals (*I*432-1), the expected Wulff polyhedral crystal habit for body-centered cubic crystals ^37^, overnight after *in vitro* mixing (Fig. 2j inset, Extended Data Fig. 6g). The crystals were further characterized by cryoEM (Fig. 2j, Extended Data Fig. 6i), SAXS (Fig. 2k) and X-ray crystallography (Fig. 2l). Indexing of the SAXS spectrum of *I*432-1 crystals indicated the *I*432 space group with a unit cell edge length of 237 Å, close to the designed lattice model (233 Å, Fig. 2k). Switching the purification tag to a chain terminus further from the crystal contact interface resulted in higher-resolution crystals (*I*432-1-CC) solved by X-ray crystallography at 3.66 Å in the designed *I*432 space group (Fig. 2l). The all-atom RMSD between the crystal structure and the computational design model was 1.34 Å over the full dihedral interface (Extended Data Fig. 7h) and 4.08 Å over two complete neighboring cages (Extended Data Fig. 7i). In solving the designed crystal structure, we also validated the structure of the O43-2 cage, which has an all-atom RMSD to the cage design model of 1.91 Å (Extended Data Fig. 7g), providing a first example of how crystal design can help address challenges in structural biology.

## Designed Crystal Engineering

An attractive property of our designed systems is that since the atomic structure is specified computationally and then genetically encoded, the properties of the crystals can be systematically modulated by further design. As a first illustration of this, we set out to tune the unit cell parameters and thus the porosity of the *F*4_1_32-2 crystal. We varied the length of the repeating arms on the C3 component of the *F*4_1_32-2 crystal (Fig. 1b) and redesigned the resulting new crystal contact interfaces, while holding the oligomer and cage assembly interfaces fixed (Fig 3a, left and top panels, Extended Data Fig. 10, methods). We obtained three new *F*4_1_32 designs that readily formed crystals (Fig 3a, lower panels) with predicted lattice parameters ranging from 351 Å to 412 Å. The lattice parameters of the actual crystals were determined by SAXS spectra analysis (Fig 3b), and were found to be remarkably close to the design values (Fig 3a, bottom row). Further, we observed that the impact of interface residue substitutions on solubility of the components and crystallization behaviors could be systematically tuned (Supplementary Table S1, Extended Data Fig. 9, Supplementary Fig. 6, Supplementary Fig. 7), which often leads to unpredictable crystallization outcomes for screened natural crystals^35^.

Compared with crystals found by screening, our designed protein crystals have distinct crystallization habits and extraordinary physical properties, which are summarized in Extended Data Table 1. Designed crystals obtained in this study occupy a unique structural space with remarkable thermal stability. They crystallize in the highly symmetrical cubic space groups (*F*4_1_32 and *I*432, Fig. 3c) and have among the highest solvent content of any in the Protein Data Bank (PDB) (as high as 90%, 5th highest among 160 thousand recorded entries, Fig. 3c inset). Despite their high solvent content, the crystals exhibit high thermostability: *I*432-1-CC remains intact after 1 hour incubation at 95 °C and even after autoclaved at 121 °C, 13 psi for 40 mins (Fig. 3d), while crystals *F*4_1_32-1 and *F*4_1_32-2 are stable up to 65°C and 85°C, respectively (Supplementary Fig. 8, crystals found by screening usually require covalent crosslinking to achieve such stability^6^). Importantly, crystal assembly is sufficiently robust to occur in complex mixtures: the two components of *I*432-1-CC are produced at high levels when expressed separately (~0.1 mg protein per mL of *E. coli* culture), and the *I*432-1-CC crystals formed with high yield upon mixing of crude cell lysates containing the two proteins (Fig. 3e, Supplementary Fig. 9a). Pure protein preparations can then be obtained by harvesting the crystals by centrifugation and washing (Supplementary Fig. 9, b-c).

We reasoned that the high stability and large open volume of our designed crystals could enable templating of inorganic nanoparticle 3D superlattices. To investigate this, we incorporated Ni-NTA coated 5 nm gold nanoparticles with the two components of the *I*432-1-CC octahedral cage, which resulted in encapsulation of the nanoparticles within the cage (Fig. 3f, Supplementary Fig. 10). The designed crystal interfaces then guided assembly of single crystalline nanoparticle arrays (Fig. 3, g-h, Supplementary Fig. 11b). Exceptionally high-quality crystals were obtained as evidenced by sharp peaks of the collected SAXS pattern as well as observation of high angle scatterings as a result of long-range ordering (Fig. 3i). To our knowledge, the level of crystallinity has not been achieved by any other colloidal assembly approaches reported thus far^17,39–41^. When crosslinked by glutaraldehyde, these nanoparticle arrays undergo repeated drying/rehydration cycles with uniform contraction/expansion (Supplementary Fig. 12), resembling recent self-adapting colloidal crystals^39^. Our designed crystal provides a general route to patterning homogeneous macroscopic single-crystalline nanoparticle arrays.

## Conclusions

Taken together, the ability to design protein crystallization and create macroscopic 3D protein materials at high accuracy by self-assembly is a substantial advance for protein design. Key to this design success is the hierarchical design of individually orthogonal interfaces that progressively constrained the system DOFs. Our approach extends previous design successes using metals^12^, DNA^17^, electrostatic^16^, and surface aromatic substitutions^18^ to generate crystal lattices from cages by programing crystal lattices at atomic accuracy, and having the self-assembly information for crystallization genetically encoded in the primary sequence of proteins. As a new class of biomaterials, computationally designed protein crystals can be prepared and purified at industrial scale, and are stable under extreme conditions (e.g., at 95°C and in lysate). The exceptionally high porosity of the crystals with channels both within and between the cages, the tunability of the lattice dimensions and hence the inter-cage channel spacings, and the ability to trigger crystallization simply by mixing components, enable a wide range of host-guest and other applications for structure determination, inorganic nanoparticle scaffolding, drug delivery, biocatalysis and biosensing.

## Acknowledgments

We thank F. Busch and V. Wysocki at Ohio State University for support with native mass spectrometry experiments. We thank G. Hura at Lawrence Berkeley National Laboratory for support and discussions with SAXS measurement. We thank X. Zuo, T. Jun at Argonne National Laboratory for help with SAXS measurement. We thank R. M. Haynes at Pacific Northwest Cryo-EM Center for help with cryoEM data collection. We thank J. Du, S. Zhang and J. De Yoreo at University of Washington for help with crystallization measurements and discussions. We thank D. Oberthür, V. Kremling, H. Chapman at Center for Free-Electron Laser Science, Hamburg, Germany, for the investigation of crystals on Free-Electron Laser. We thank S. Weaver, K. Patel and T. Gonen at University of California, Los Angeles for help with microED. We thank S. Dickinson, N. Bethel and M. Wu at University of Washington for help with cryoEM sample preparation and screening. We thank S. Caldwell at University of Washington for help with analyzing crystallographic data. We thank X. Wang at University of Washington for suggestions on X-ray crystallography data collection. We thank T. Huddy, R. Kibler, N. Bethel, A. Khmelinskaia, D. Zambrano and R. Haas at the University of Washington for providing potential protein building blocks. We thank H. Bai, R. Kibler, T. Huddy, H. Pyles, C. Xu and A. Ljubetic at the University of Washington for help with scripting and softwares. We would also like to thank members of Baker lab and Institute for Protein Design, particularly J. Bale, N. King and F. Dimaio, for useful discussions.

## Author contributions

Conceptualization: U.N., Z.L., S.W., E.C.Y. and D.B.; Methodology: Z.L., S.W., U.N., W.S. and D.B.; Investigation: Z.L., S.W., U.N., E.C.Y., R.D. and M.M.; CryoEM: Z.L., S.W., A.J.B.; X-ray Crystallography: Z.L., S.W., U.N., A.K.B., M.J.B., H.N., A.K. and B.S.; SAXS: S.W., B.L. and S.S.; Building blocks: Z.L., U.N., E.C.Y., J.M.L., Y.H., Q.D., A.F., N.I.E. and W.Y.; Design protocols: Z.L., S.W., U.N., W.S., Y.H., A.C., Q.D. and A.E.; Rosetta Scorefunction: H.H.; Visualization: Z.L., S.W., U.N., A.J.B., W.Y. and D.B.; Funding acquisition: D.B.; Supervision: D.B.; Writing – original draft: Z.L., S.W., U.N. and D.B.; Writing – review and editing: Z.L., S.W., U.N., A.K.B., A.J.B., E.C.Y., H.H. and D.B.

## Methods

### Cage docking and design

A library of cyclic oligomer scaffolds (C2, C3, C4) from crystal structures deposited in the PDB (http://www.rcsb.org/pdb/) and from previous *de novo* designs^28,30,35,42^ were docked into tetrahedral and octahedral cages (T32, T33, O32, O42 and O43). Cage dockings were carried out by RPXDock, which used hierarchical sampling of residue pair transform scoring to find high designability docking^32^. The top 100 to 500 dockings of each symmetry were sequence designed by Rosetta^24^ using a protocol based on two-component protein-protein interface design methods implemented within the RosettaScripts framework^43^. Either beta_nov16 or a clash-fixed score function was used during the design. The protein-protein interfaces were designed with rigid protein backbones by packing rotamers with layer design restrictions at interacting residues. All cage designs were filtered by a number of metrics evaluating the designed cage interface, including methionine count <=5, shape complementarity >0.6, ddG <-20 kcal/mol, sasa (solvent-accessible surface area) <1,600, clash check <=2, and unsatisfied hydrogen bonds <=2, before visual inspection of the hydrophobic packing for more than two pairs of intersected hydrophobic sidechains. A final filtering by placing the cages into the corresponding Wyckoff sites in the target crystal lattice and inspecting the cage backbone for sufficient secondary structure interaction between the cages (Extended Data Fig. 5, before the extraction of dihedrals). Cage designs with no or little interacting secondary structures were left out for experimental verification.

### Crystal docking and design

Ten previously designed cages (T3-10, T32-28, T33-09, T33-15, T33-21, T33-23, T33-28, T33-31, T33-51 and T33-53) ^29,31,33,34^ and experimentally verified new cages were used for the docking to target the *P*23, *P*432, *P*4_2_32, *I*432, *F*23, *F*432, and *F*4_1_32 space groups. Dockings were performed in Pymol^44^ by alignment and translations. For the docking of the *F*4_1_32 space group, two tetrahedral cages (TET-1, TET-2) were aligned with every C2 symmetry axis along the x,y, and z coordinate axes and center of mass at the origin of the coordinate (Extended Data Fig. 5a, left panel). While keeping TET-1 fixed at the origin, TET-2 was rotated along the z-axis for 90 °, and translated along (1,1,-1) direction to dock with TET-1 (until distance between the backbone of the two cages was less than 6 Å with 1 Å increment translations, Extended Data Fig. 5a, middle panel). The hexamer between the two cages of D3 dihedral symmetry was extracted and aligned with its C3 axis along the z-axis, C2 axis along the x-axis, and center of mass at origin (Extended Data Fig. 5a, right panel). The docking of the *I*432 space group was similar between two octahedral cages (OCT-1, OCT-2), except OCT-2 was only translated without any rotation (Extended Data Fig. 5b). In both cases, the D3 assembly units contained the crystal contact and were used for Rosetta sequence design. In the RosettaScripts framework, input monomers were symmetrized into D3 dihedrals, and then we sampled the interface distances by translation along the z-axis (dihedral axis) within ± 1.5 Å of the docked conformation without rotation DOF. For each translation, the dihedral interface was designed with rigid protein backbones by packing rotamer with layer design restrictions at interacting residues. Then designs were filtered by ddG <0, sasa >200, clash check <=2, and unsatisfied hydrogen bonds <=2 before being visually inspected for hydrophobic packings. Working crystal designs usually have one to three pairs of intersected hydrophobic sidechains at each C2 branch of the D3 dihedral. For DOF of each design step: two cyclic oligomers have eight: two monomers, each has four DOFs into cyclic oligomers^28^, co-assembly of the cage has four: two cyclic oligomers, each has two DOFs into cage^29^, and the final crystal has only one: one cage, only one DOF.

### Redesign of crystal unit cell dimension

To design the unit cell dimension of the *F*4_1_32-2 crystals, the DHRs at the periphery of the cage were engineered with new fusions to alter the cage spacing in the crystal lattice. From the cryoEM model of the T32-15 crystal, the C3 symmetry unit was extracted (Extended Data Fig. 10a) and fused with a library of verified DHRs by WORMS protocol^42^ (fusion region defined as residue 1-195, away from the oligomer and cage assembly interfaces, Extended Data Fig. 10b). The fusions with new backbone were docked by RPXDock^32^ in D3 symmetry with only translation DOF along the C3 axes of the dihedral (Extended Data Fig. 10c). The docking angle for crystal propagation was pre-implemented before C3 extraction, and fusions showing no contact by docking were filtered from downstream design. The monomer of the docked dihedral was extracted and designed at the fusion region by Rosetta by packing rotamers with layer design restrictions and filtered by shape complementarity >0.5, alanine percentage < 40% at the junction (Extended Data Fig. 10d). The fusion designs were then filtered with AlphaFold2 predictions (model 4, RMSD<2 design model vs prediction, pLDDT>85, Extended Data Fig. 10e). As the final design step, filtered fusions were Rosetta designed in D3 symmetry by the same protocol as described above for crystal contacts (Extended Data Fig. 10f). Both the designs of the fusion junction and the crystal contacts were experimentally verified at the same time by crystal assembly (Extended Data Fig. 10g).

### Rotation correction for design /crystal model comparison

Because of the minimization of the virtual connections (“jumps”) between the symmetry subunits in Rosetta design, the D3 dihedral design outputs could slightly deviate from the restricted rotation constrained by crystal symmetry. In order to construct a design model of the crystal unit cell to compare with the crystallographic model, rotation-corrected dihedral models were prepared by applying the same dihedral axis displacement as in the design model, while keeping the rotation angle fixed (before minimization). The rotation-corrected dihedral models were then used to propagate the design model of the cage into one crystal unit cell by alignment.

### Clash-fixed Rosetta score function

We observed that designs made with the beta_nov16 score function tend to have higher levels of steric clashing than observed in high-resolution crystal structures of native proteins. Parameters in the Rosetta score function was refitted to reduce this steric-clashing propensity, resulting in the “clash-fixed” score function. We are preparing a manuscript that describes this problem and the new score function in more detail. The files in the “clash-fixed_scorefunction” folder of supplementary material encode all parameter changes and can be used to implement the new score function in design.

### Source code, examples, and design models

Examples of design scripts, symmetry definition files, and design models (cages including T32-15, T32-15-6H, O43-2, T33-ECY54, T33-ECY55, T33-ECY59, T33-ECY66, T33-ECY67, O43-UWN38, O43-UWN453, O43-ZL1, O43-ZL7, O32-ZL4, O43-EK1, and crystal contact dihedrals of F4_1_32-1-0, F4_1_32-2, I4_1_32-1, F4_1_32-2-ex1, F4_1_32-2-ex2 and F4_1_32-2-ex3) can be found in the supplementary material.

### Protein expression and purification

Synthetic genes were optimized for *E. coli* expression and purchased from IDT (Integrated DNA Technologies) as plasmids in pET29b vector with a hexahistidine affinity tag. For cage screening, genes for the two cage components were joined together by another RBD domain (gene sequence: TAAAGAAGGAGATATCATATG) in between, and the hexahistidine tag was added on only one of the components. Plasmids were cloned into BL21* (DE3) *E. coli* competent cells (Invitrogen). Single colonies from agar plate with 100 mg/L kanamycin were inoculated in 50 mL of Studier autoinduction media ^45^, and the expression continued at 37 °C for over 24 hours. The cells were harvested by centrifugation at 4000 g for 10 min, and resuspended in 35 mL lysis buffer of 300 mM NaCl, 25 mM Tris pH 8.0 and 1 mM PMSF. After lysis by sonication and centrifugation at 14000 g for 45 min, the supernatant was purified by Ni^2+^ immobilized metal affinity chromatography (IMAC) with Ni-NTA Superflow resins (Qiagen). Resins with bound cell lysate were washed with 10 mL (bed volume 1 mL) of washing buffer (300 mM NaCl, 25 mM Tris pH 8.0, 60 mM imidazole) and eluted with 5 mL of elution buffer (300 mM NaCl, 25 mM Tris pH 8.0, 300 mM imidazole). Both soluble fractions and full cell culture were checked by SDS-PAGE. Soluble designs with correct molecular weight and cage designs with pull-down showing two separate bands were further purified by size exclusion chromatography (SEC). Concentrated samples were run in 150 mM NaCl, 25 mM Tris pH 8.0 on a Superose 6 Increase 10/300 gel filtration column (Cytiva). SEC-purified designs were concentrated by 10K concentrators (Amicon) and quantified by UV absorbance at 280 nm before further assembly and characterization.

### Crystallization of designs

For all crystallization experiments, the concentration of protein monomers was used for simplification. For example, “50 μM component A” represents 50 uM of component A monomer, thus 50/n μM of oligomer depending on the n fold of A (n = 2, 3, 4); “50 uM cage” represents cages assembled from 50 uM component A and 50 μM of component B, thus 50/12 = 4.2 μM cage assembly for tetrahedral cages (12 copies of each component), and 50/24 = 2.1 μM cage assembly for octahedral cages (24 copies of each component).

Since our crystals were designed to crystallize with no dependence on specific precipitants or additives, all crystallization experiments were screened in NaCl solution. As initial tests, cage components were mixed in 150 mM NaCl, 25 mM Tris pH 8.0 from overnight to days to observe cage or crystal assembly. For designs that didn’t crystallize, the mixed solution was set up for hanging drops with 0.5 M to 5 M NaCl. For each screened NaCl concentration, NaCl solutions of the target concentration were used as reservoir solution, and the hanging drop was a mixture of 2 μL of the solution of the mixed components and 2 μL of the reservoir solution. For some designs like *I*432-1-CC, the hanging drop was set up with only the cage solution (no extra NaCl) against NaCl reservoir solution. Crystal trays were checked for crystallization for up to two weeks and further optimized with NaCl concentration. Designs were also examined for batch crystallization by mixing cages and NaCl solution at 1:1 volumetric ratio according to the hanging drop conditions. Crystallization conditions for all designs were summarized in Supplementary Table S1.

### Thermal stability of designed protein crystals

*F*4_1_32-1 crystals by mixing, *F*4_1_32-2 crystals by hanging drop (diluted in 1.5 M NaCl), and *I*432-1-CC crystals by hanging drop (diluted in 0.5 M NaCl) were used for thermal stability study. 10~20 μL of the crystals were intubated in T100 Thermal Cycler (Bio-Rad) at 55 °C, 65 °C, 75 °C, 85 °C and 95 °C for 1 hour. Crystals were checked before and immediately after incubation (within 5 mins) under the optical microscope by pipetting the solution and applying to a siliconized glass slide. Autoclaving experiment was done for 20 μL of crystal solution in an opened 1.5 mL Eppendorf tube at 121 °C, 13 psi for 40 mins.

### Purification of *I*432-1-CC crystals from lysate

Clarified lysate after sonication and centrifugation (see protein expression and purification section) of the two components for *I*432-1-CC crystals were mixed at 1:1 volumetric ratio. 20-50 μm of crystals appeared in solution after overnight incubation. The crystals were collected by centrifugation of 4000 g for 10 min, and washed twice with 300 mM NaCl, 25 mM Tris pH 8.0. For recrystallization, the centrifuged crystals were dissolved in water, and set up as hanging drops in 0.5 M NaCl. Despite the high thermal stability of designed crystals, recrystallization demonstrated that the crystal assembly is reversible by controlling ionic strength, providing flexibility for storage and applications.

### Scaffolding 3D gold nanoparticle arrays

The constituent cage of *I*432-1-CC crystal has an interior void of ~10 nm diameter, and the His-tags from both components facing towards the inside of the cages provide sites for host-guest interaction. We used the Ni-NTA His-tag interaction (K_D_≈10^−13^ M at pH 8.0)^46^ to guide the encapsulation of 5 nm AuNPs into the interior of the O43 cages. 50 μM *I*432-CC-1 cages were assembled overnight in 150 mM NaCl, 25 mM Tris pH 8.0. Then 10 μL cage, 2 μL 1 M imidazole and 88 μL of 5 nm gold nanoparticles (AuNPs) functionalized with nickel (II) nitriloacetic acid (NTA) chelates (0.5 μM in 50 mM MOPS, pH 7.9, Nanoprobes) were mixed for nanoparticle encapsulation. Because of the low ionic strength (~15 mM NaCl) and excess amount of AuNPs (>4 times excess), the cages will have more transient self-assembly to have oligomer pieces disassembled and assembled^47^, giving AuNPs time to bind with partially disassembled cage and be encapsulated. The encapsulation was characterized by nsEM by diluting the solution 10 times in 150 mM NaCl, 25 mM Tris pH 8.0, 300 mM imidazole right before imaging. It took about two weeks for more than half of the cages to have AuNPs encapsulated. For purification, 15 μL of 1 M NaCl and 30 μL of 1 M imidazole were added to the solution right before centrifugation at 14000 g for 30 mins, the pellets of free AuNPs and AuNPs encapsulated in cages were redispersed by 100 μL 150 mM NaCl, 25 mM Tris pH 8.0, 300 mM imidazole and centrifuged again at 14000 g for 30 mins. The final pellet was dispersed in 20 μL of 150 mM NaCl, 25 mM Tris pH 8.0, 300 mM imidazole. The solution, now containing O43 cages with AuNP encapsulation and free AuNPs (which didn’t assemble with the cage), was set up for crystallization by hanging drop of 4 μL solution against 0.5 M NaCl. We also found that cages of *I*432-1 crystals with His-tags facing towards the outside of the cage, co-assembled with the AuNPs into polycrystalline binary lattices upon mixing with the AuNPs (Supplementary Fig. 11a).

### Crystals structure determination

Data collection was performed with synchrotron radiation either at the Advanced Photon Source (APS) on beamline 24ID-C or at the Advanced Light Source (ALS) on beamline 8.2.1/8.2.2. X-ray intensities and data reduction were evaluated and integrated using either XDS^48^ or HKL3000^49^ and merged/scaled using Pointless/Aimless in the CCP4 program suite^50^. Structure determination and refinement starting phases were obtained by molecular replacement using Phaser^51^ using the design model for the structures. Following molecular replacement, the models were improved using phenix.autobuild^52^; efforts were made to reduce model bias by setting rebuild-in-place to false, and using simulated annealing and prime-and-switch phasing. Structures were refined in Phenix^52^. Model building was performed using COOT^53^. The final model was evaluated using MolProbity^54^. Data collection and refinement statistics are recorded in Table S2. The final structure was deposited in the Protein Data Bank (PDB), http://www.rcsb.org/, under PDB accession number 8CUS, 8CUT, 8CUU, 8CUV, 8CUW, 8CUX, 8CWS, 8CWZ.

### Transmission negative-stain electron microscopy (nsEM)

Cage fractions from SEC traces or cages by *in-vitro* mixing were diluted to 0.5 μM (monomeric component concentration) for negative-stain EM characterization. For the crystal samples, crystals were first crushed into smaller pieces under the optical microscope by cryo-loops before applying onto the EM grids. A drop of 6 μL sample was applied on negatively glow discharged, formvar/carbon supported 400-mesh copper grids (Ted Pella, Inc.) for more than 2 mins. The grid was blotted and stained with 3 μL of uranyl formate, blotted again, and stained with another 3 μL of uranyl formate for 20 s before final blotting. 0.75% and 2% uranyl formate were used for different samples. The screening was performed on either a 120kV Talos L120C transmission electron microscope (Thermo Scientific) or a 100kV Morgagni M268 transmission electron microscope (FEI).

### NsEM image processing

All nsEM datasets were processed by CryoSparc software^55^. Micrographs were imported into the CryoSparc web server, and the contrast transfer function (CTF) was corrected. Around 100 particles were manually picked, 2D classified and selected classes were used as templates for particle picking in all images. All the picked particles were 2D classified for 20 iterations into 50 classes. Particles from selected classes were used for building the ab-initio initial model. The initial model was homogeneously refined using C1 and the corresponding T/O symmetry.

### CryoEM sample preparation

For the T32-15 cage, 2 μL of 5.3 mg/mL of T32-15 cage in 150 mM NaCl, 25 mM Tris pH 8.0 was applied to glow-discharged C-flat holey carbon grids. Vitrification was performed via manual blotting at ambient temperature and humidity. For the crystal samples, crystals were first crushed into smaller pieces under the optical microscope by cryo-loops right before freezing. Samples in the corresponding crystallization buffer were frozen using glow-discharged 1.2/1.3-T C-flat holey carbon grids and a Mark IV Vitrobot with a wait time of 5 seconds, a blot time of 7.5 seconds, and a blot force of 0 before being immediately plunge frozen into liquid ethane.

### CryoEM data collection

T32-15 data collection was performed automatically using Leginon^56^ to control a ThermoFisher Titan Krios 300 kV TEM (PNCC Krios 1) equipped with a standalone K3 Summit direct electron detector^57^ and operating in super-resolution mode. Random defocus ranges spanned between −0.8 and −2.2 μm using image shift, with one-shot per hole, and 9 holes per stage move. 4202 movies were collected with a pixel size of 0.5144 Å with a total dose of 50 e^-^/A^2^. Individual images of crushed crystals were manually collected on either a ThermoFisher Glacios 200 kV TEM equipped with a K2 summit direct electron detector using serialEM, or a ThermoFisher Talos L120C TEM equipped with a BM-Ceta camera using EPU 2.0.

### CryoEM data processing

All data processing was carried out in CryoSPARC^55^. Alignment of movie frames was performed using Patch Motion with an estimated B-factor of 500 Å^2^, maximum alignment resolution set to 3. Outputs were binned to a final pixel size of 1.0288 Å/pixel by setting the output F-crop factor to ½. Defocus and astigmatism values were estimated using Patch CTF with default parameters. 727,160 particles were picked in a reference-free manner using Blob Picker and extracted with a box size of 320 Å. An initial round of reference-free 2D classification was performed in CryoSPARC using 100 classes and a maximum alignment resolution of 6 Å. The best classes - composed of 207,495 particles - were then used for 3D ab initio determination using the C1 symmetry operator. These same 2D classes were next low pass filtered to 20 Å and used as templates for a second round of particle picking using Template Picker, resulting in a new set of 964,346 particle picks which was extracted with a box size of 320 pixels. Following another round of reference-free 2D classification, the best 511,464 particles were submitted for non-uniform refinement in the presence of T symmetry for a final estimated global resolution of 3.34 Å. Local resolution estimates were in CryoSPARC using an FSC threshold of 0.143. 3D maps for the two half maps, final unsharpened map, and the final sharpened map were deposited in the EMDB under accession number EMD-27031.

### CryoEM model building and validation

The design mode of the T32-15 cage was used as an initial reference for building the final cryoEM structure. The model was manually edited and trimmed using Coot ^53,58^. We then further refined the structure in Rosetta using density-guided protocols^59^. This process was repeated iteratively until convergence and high agreement with the map was achieved. Multiple rounds of relaxation and minimization were performed on the complete cage and were manually inspected for errors each time. Throughout this process, we applied strict non-crystallographic symmetry constraints in Rosetta^60^. Residues 1-124 for each chain were truncated to the cβ carbon due to the low resolution of this region and low confidence in determination of sidechain rotamers. Phenix real-space refinement was subsequently performed as a final step before the final model quality was analyzed using Molprobity^54^ and EM ringer^61^. Figures were generated using either UCSF Chimera^62^ or UCSF ChimeraX^63^. The final structure was deposited under PDB accession number 8CWY.

### Small-angle X-ray scattering (SAXS) of crystals

All crystal structures in this work were confirmed primarily by SAXS. SAXS characterization was carried out at the 12-ID-B beamline of the Advanced Photon Source at Argonne National Laboratory following a literature report^64^. The wavelengths of X-rays used at 12 was 0.9322 Å (13.3 keV), and the system was calibrated using silver behenate as a standard. Two sets of slits were used to define and collimate the X-ray beam, and parasitic scattering was removed using a pinhole. Typical exposure times of between 0.1 and 0.5 s were used. The scattered X-rays were collected with a charge-coupled device (CCD) detector, and one-dimensional scattering profiles were obtained by radially averaging the images into scattering intensities, *I*(*q*), against the scattering vector magnitude, *q*. In this work, all *I*(*q*) profiles were plotted on a logarithmic scale for clarity.

### Modeling of protein crystal SAXS data

One-dimensional SAXS profiles were simulated and indexed following a literature report ^65,66^. Crystallographic symmetries and lattice parameters are determined by indexing powder diffraction patterns. The full-atom representation of protein cages is located into their corresponding Wyckoff positions of the lattice, from the smallest multiplicity site to the higher. The average crystalline domain size, Debye–Waller factor (DWF) and microstrain parameter were empirically adjusted to fit the peak width and relative peak intensities across the entire pattern.

In Supplementary Fig. 13, the calculated form factor of each cage is shown in blue and intensity profiles are shown in red and black for simulated and measured profiles, respectively. For all three curves, simulated peaks at around the form factor minima deviate from the measured ones than other peaks. This might indicate that the form factor minima in the real samples are not as sharp as the ones calculated or might have been shifted slightly. This is possible when there is a size distribution of the cages, for example due to different hydration of the cages, or when there are diffuse scatterings due to orientational or positional defects. We do not neglect non-perfect random orientation of the crystals in solution, which will cause a certain random peak stronger than the other.

---

**Extended Data Figure 1.**
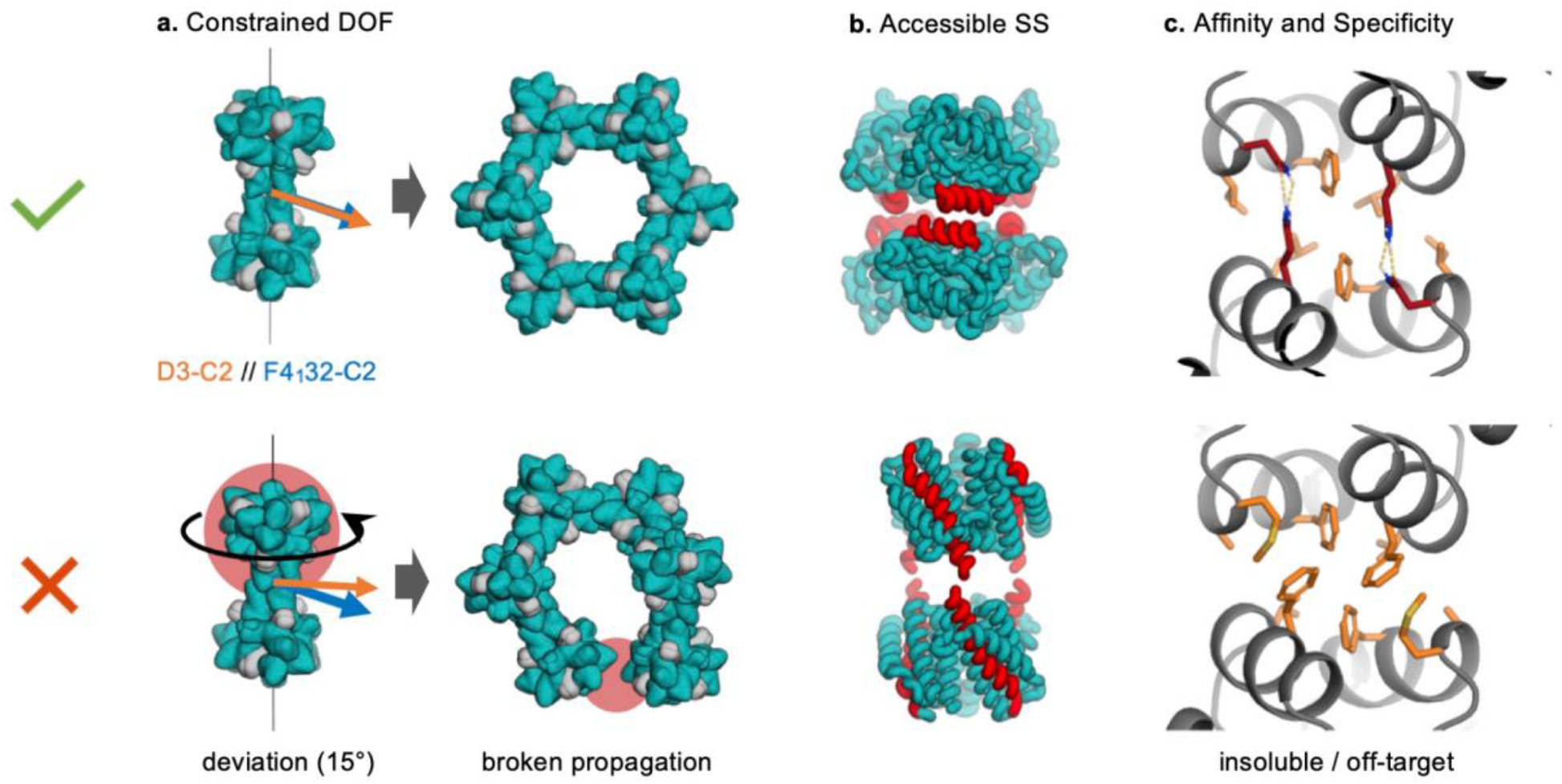
Design rules of 3D protein crystals. **a**, Constrained degree of freedom (DOF): The angle of rotation at the designed dihedral crystal interface (Fig. 1a, right panel) must be precisely specified by the design process, where the C2 axis of the dihedral needs to coincide with the C2 axis of the space group. In this example, the disruptive effect (highlighted in red) of a 15-degree error in alignment on crystal assembly is illustrated; similar crystal lattice breakdowns occur with all deviations from the target alignment angle. **b**, Accessible secondary structure (SS): Dihedral interfaces with helices perpendicular to the symmetry axis (docked from T33-15 cage) are more designable than those with helices parallel to the symmetry axis (docked from T33-21 cage^29^). Interacting secondary structures are highlighted in red. **c**, Affinity and Specificity: Working interfaces have sufficient hydrophobic packing with specific polar interactions at the boundary. Highly hydrophobic interfaces destruct the designed self-assembly, including insoluble components and off-target assemblies.

**Extended Data Figure 2.**
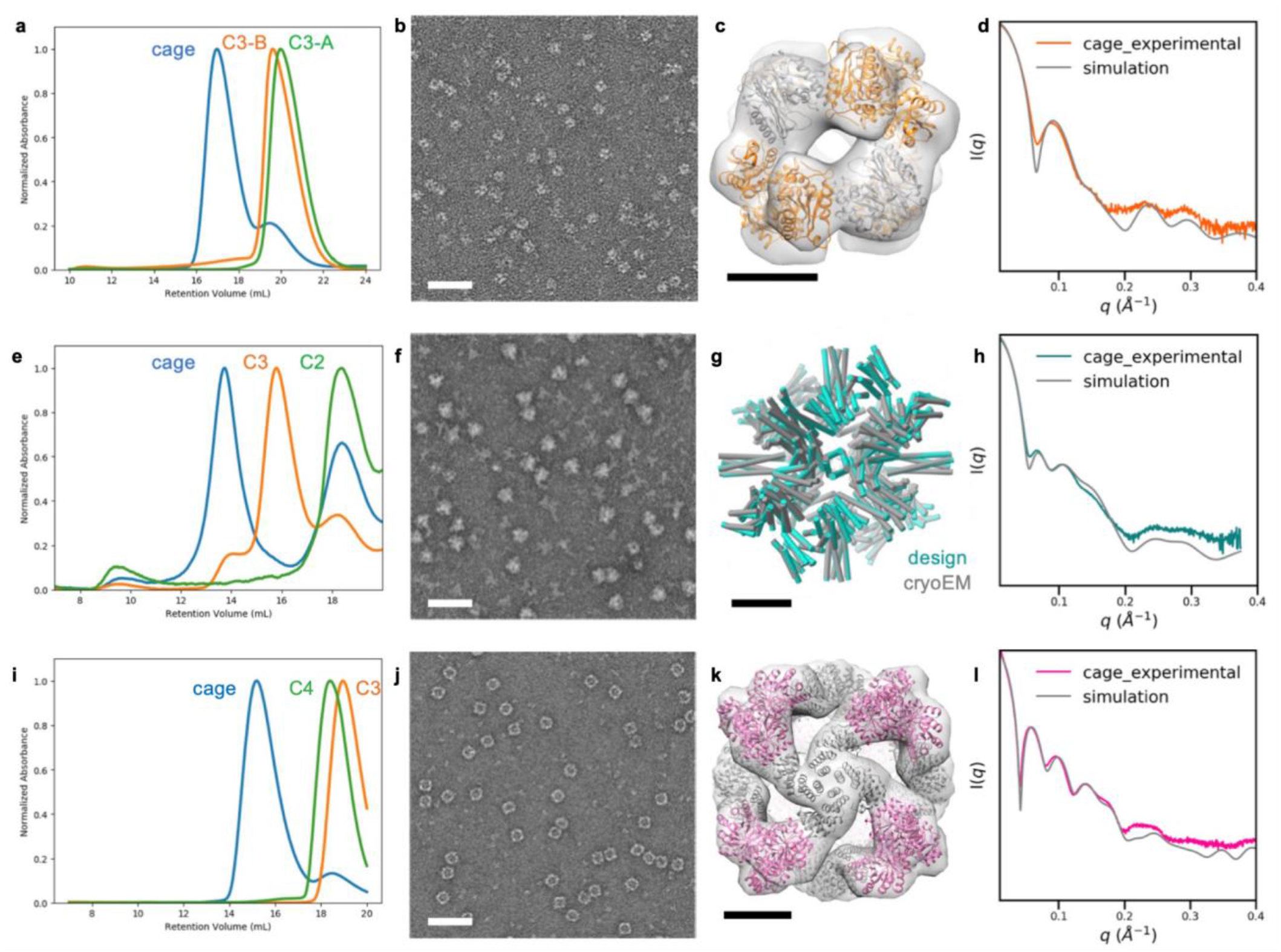
Characterizations of the constituent cages of designed crystals. **a-d**, T33-15-D3-4, **e-h**, T32-15, **i-l**, O43-2. **a**,**e**,**i**, SEC chromatograms of two oligomeric components (green and orange) and cages assembled via *in-vitro* mixing of components (blue). **b**,**f**,**j**, nsEM images (scale bars, 50 nm). **c**,**g**,**k**, overlay of the design model with 3D reconstructed nsEM density map/ cryoEM model (scale bars, 5 nm). **d**,**h**,**l**, SAXS profile and simulation results of cages.

**Extended Data Figure 3.**
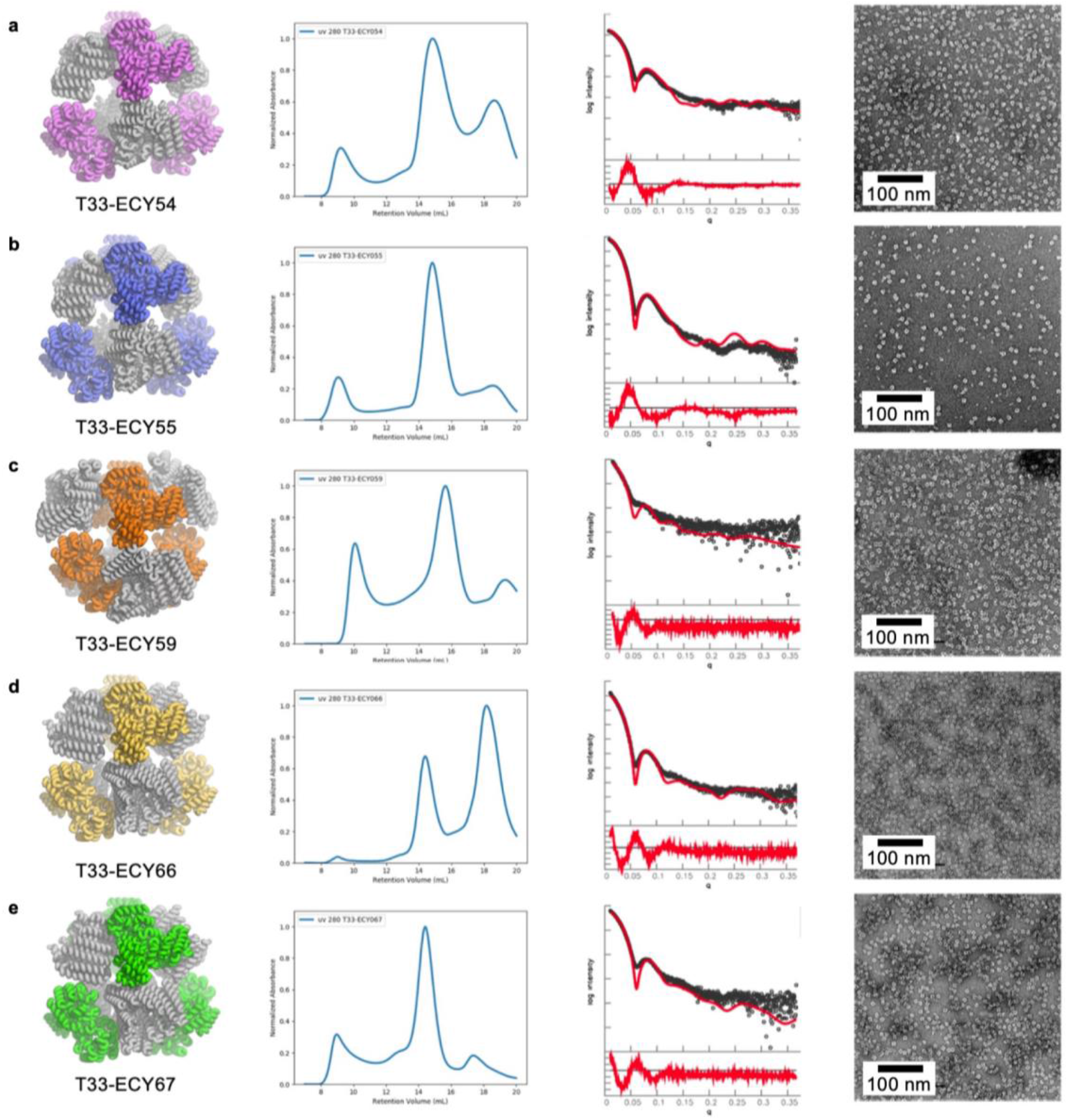
Characterizations of new tetrahedral cages for crystal design. **a-e**, from left to right, computational model, SEC chromatogram, SAXS profile, and nsEM images.

**Extended Data Figure 4.**
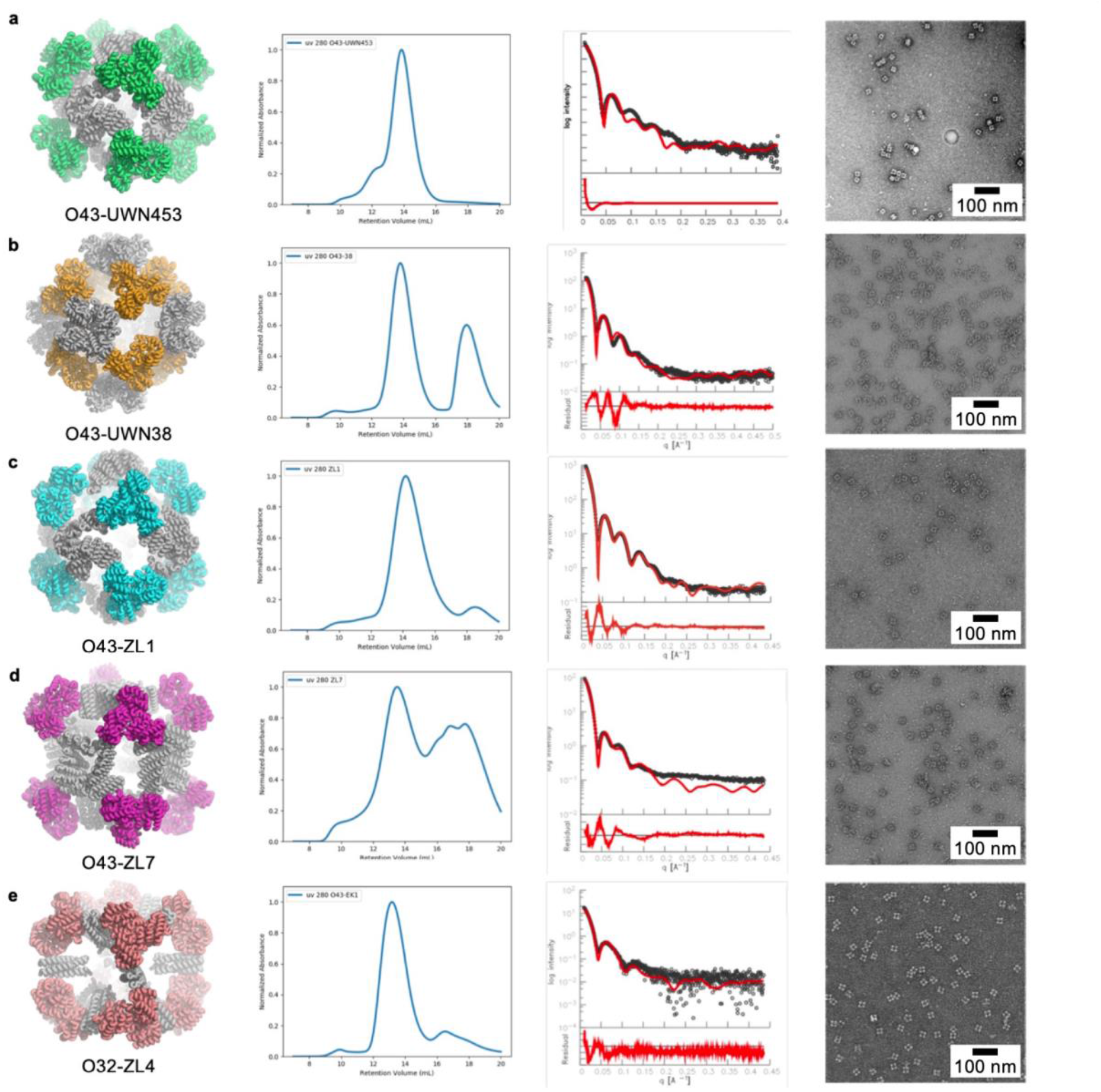
Characterizations of new octahedral cages for crystal design. **a-e**, from left to right, computational model, SEC chromatogram, SAXS profile, and nsEM images.

**Extended Data Figure 5.**
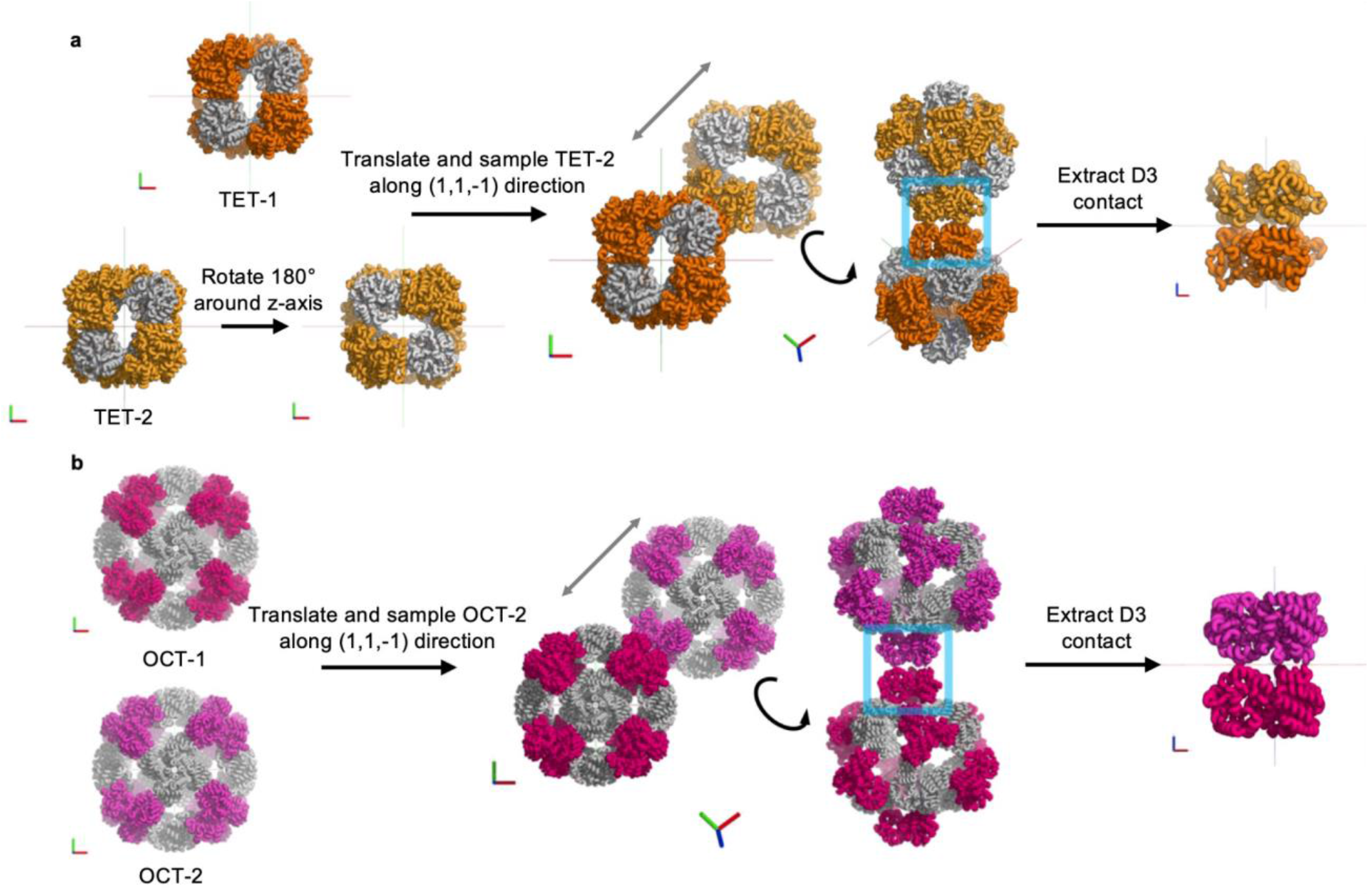
Symmetric dockings of tetrahedral and octahedral cages into crystal lattices. **a**, Two tetrahedral cages are docked along their C3 axis for crystal contacts of D3 dihedrals, which allow them to crystallize in the *F*4_1_32 space group. **b**, Two octahedral cages are docked along their C3 axis for crystal contacts of D3 dihedrals, which allow them to crystallize in the *I*432 space group. See methods for detailed docking protocol.

**Extended Data Figure 6.**
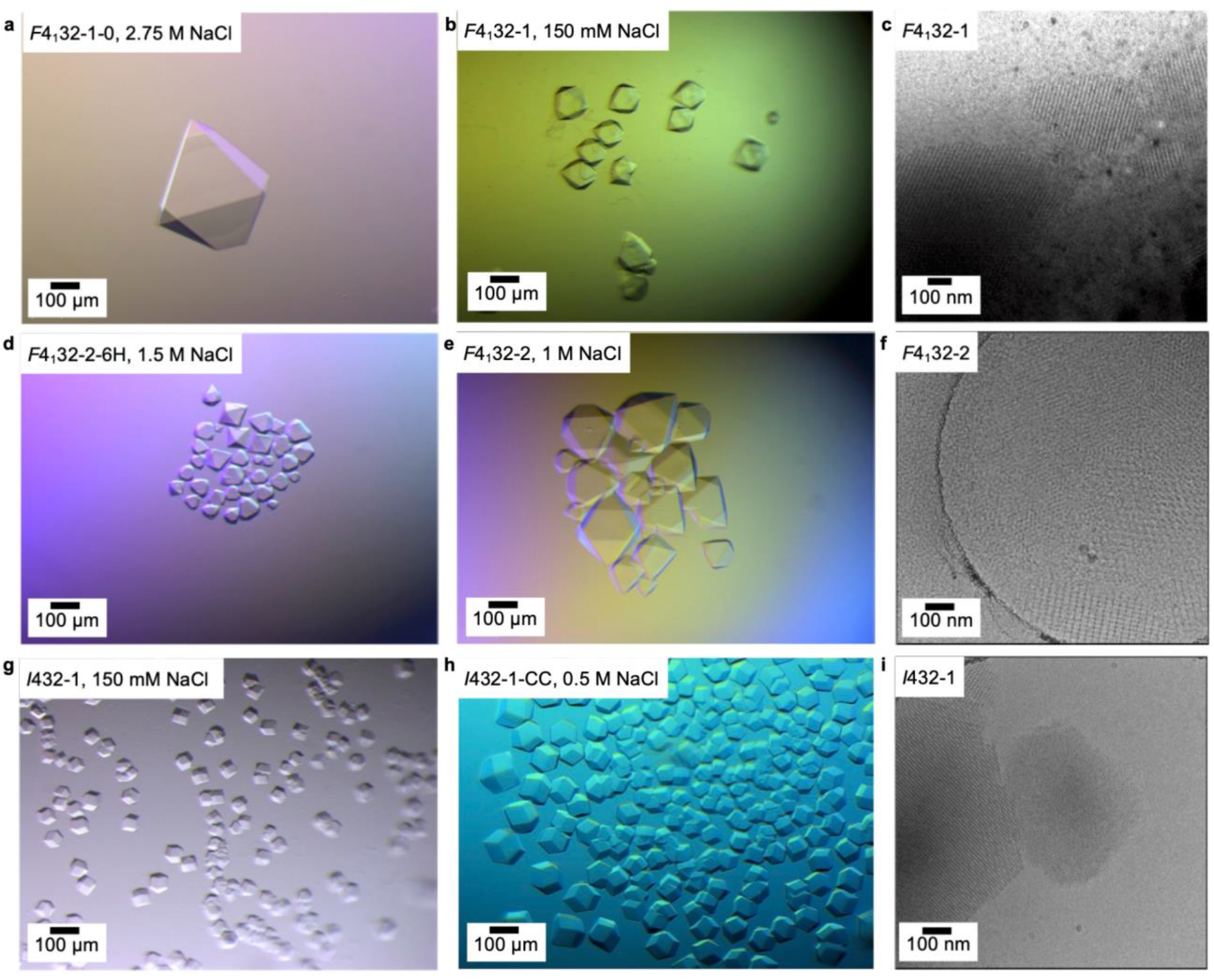
Optical microscopy and cryoEM characterization of designed protein crystals. **a**, Optical micrograph of *F*4_1_32-1-0 crystals. **b**, Optical micrograph of *F*4_1_32-1 crystals. **c**, CryoEM image of *F*4_1_32-1 crystals. **d**, Optical micrograph of *F*4_1_32-2-6H crystals. **e**, Optical micrograph of *F*4_1_32-2 crystals. **f**, CryoEM image of *F*4_1_32-2 crystals. **g**, Optical micrograph of *I*432-1 crystals. **h**, Optical micrograph of *I*432-1-CC crystals. **i**, CryoEM image of *I*432-1 crystals.

**Extended Data Figure 7.**
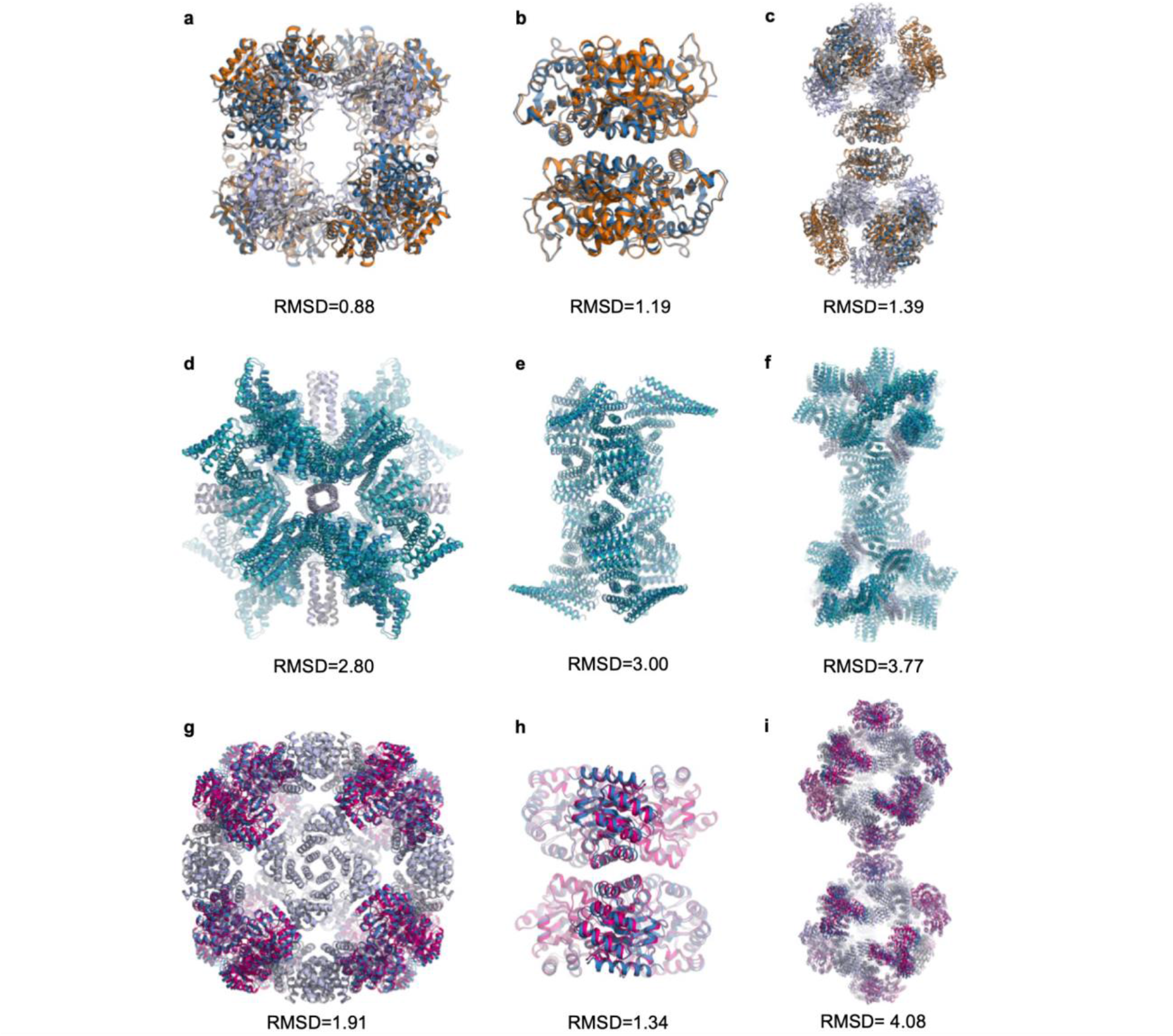
The alignment of X-ray structure and computational design model (orange, teal, hot pink and gray) of designed protein crystals (sky blue and light blue). **a**, Alignment of T33 cages of *F*4_1_32-1 crystals. **b**, Alignment of D3 dihedrals of *F*4_1_32-1 crystals. **c**, Alignment of two neighboring cages of *F*4_1_32-1 crystals. **d**, Alignment of T32 cages of *F*4_1_32-2 crystals. **e**, Alignment of D3 dihedrals of *F*4_1_32-2 crystals. **f**, Alignment of two neighboring cages of *F*4_1_32-2 crystals. **g**, Alignment of O43 cages of *I*432-1 crystals. **h**, Alignment of D3 dihedrals of *I*432-1-CC crystals. **i**, Alignment of two neighboring cages of *I*432-1-CC crystals. All-atom RMSDs are shown below each alignment.

**Extended Data Figure 8.**
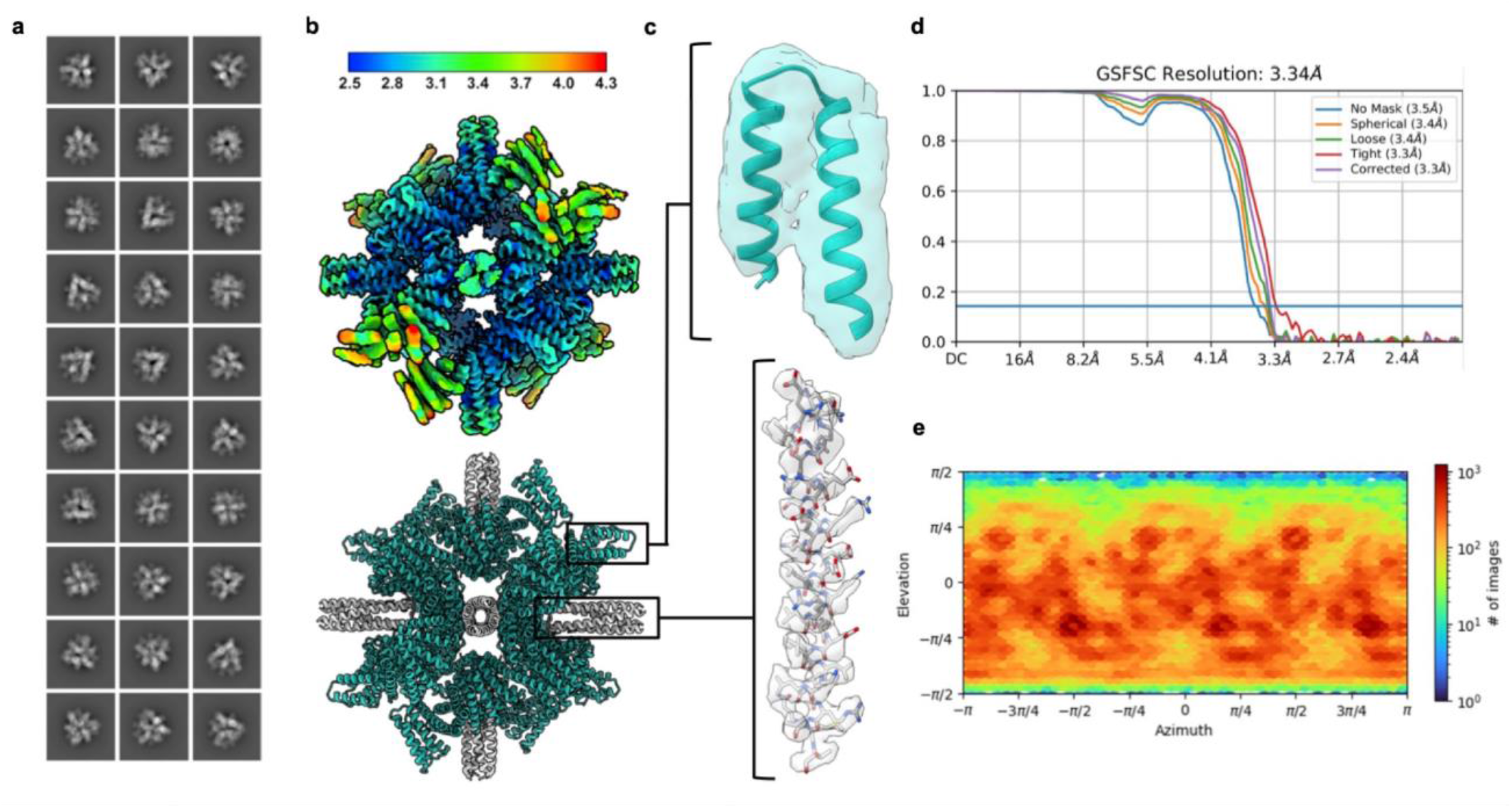
CryoEM data of the T32-15 cage. **a**, Representative 2D class averages of the T32-15 cage. **b**, CryoEM local resolution map of the T32-15 cage (top) and built atomic model (bottom). Local resolution estimates range from ~2.5 Å at the core to ~4 Å along the crystal-contact forming helices. **c**, Map-to-model comparison within a low-resolution region (top) and a high-resolution region (bottom). **d**, Global FSC. **e**, Orientational distribution plot demonstrating full angular sampling.

**Extended Data Figure 9.**
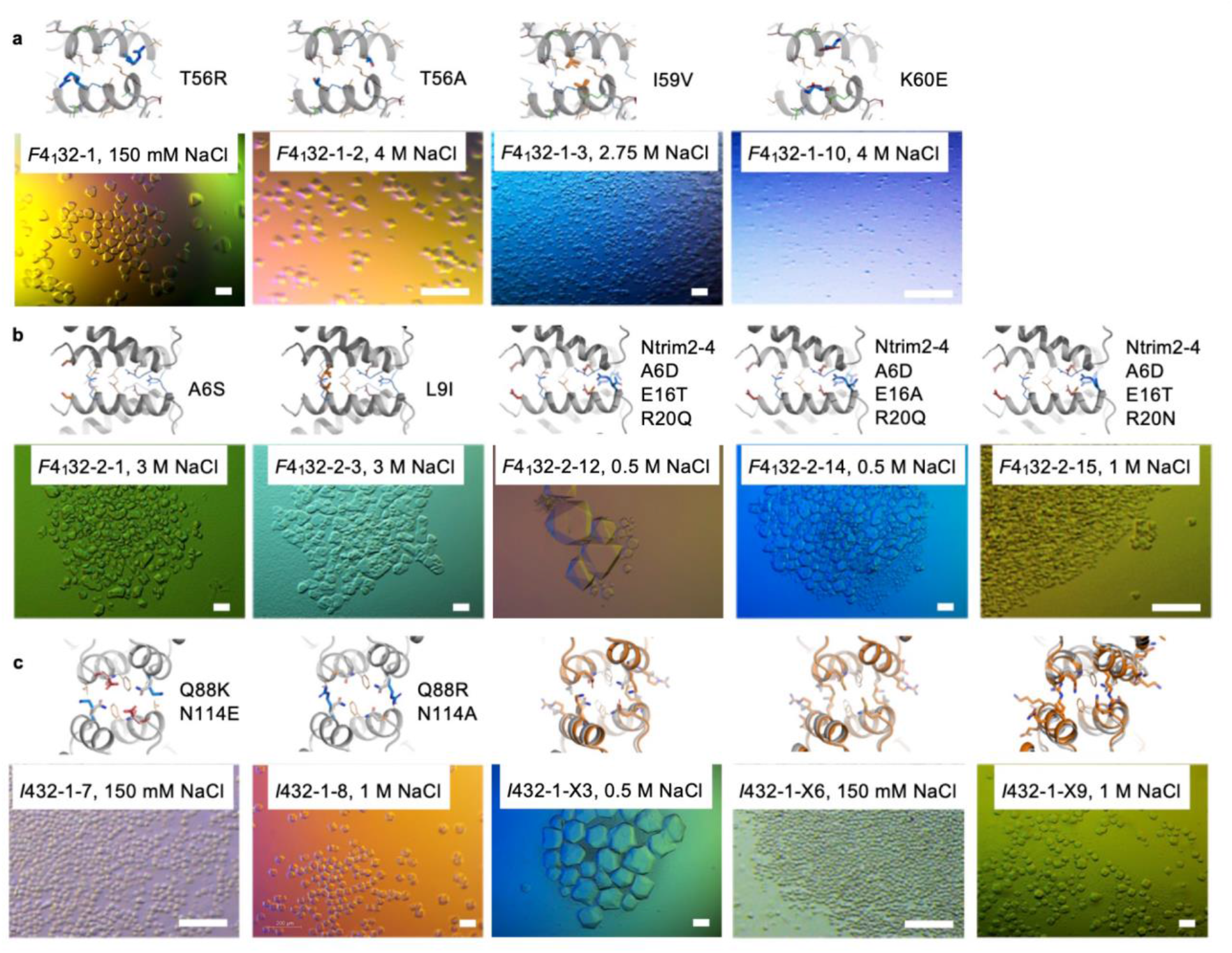
Tuning the crystallization behavior of designed crystals by mutagenesis. **a**, Mutations to the *F*4_1_32-1 crystals. **b**, Mutations of *F*4_1_32-2 crystals. **c**, Mutations and redesigns (orange) of *I*432-1 crystals. Top panels, crystal interface models based on X-ray structure. Interface side chains are hypothetically placed to demonstrate mutation sites. Bottom panels: optical micrographs of representative crystallization results. Scale bars, 100 μm.

**Extended Data Figure 10.**
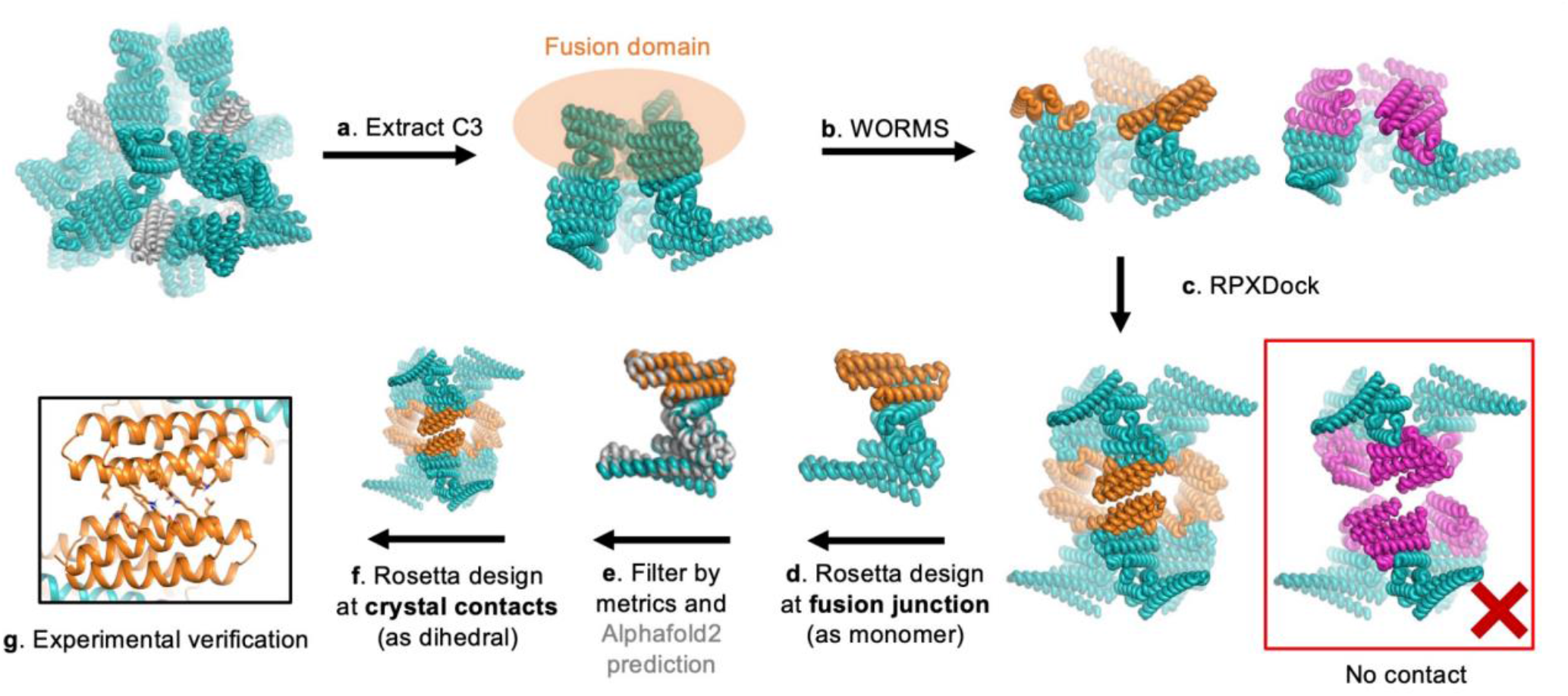
Design pipeline for engineering crystal unit cell dimension. The crystal contact of the *F*4_1_32-2 crystal was redesigned with different DHR arm fusion. See methods for the details of step **a-g**.

**Extended Data Table 1.** Comparison of properties between designed protein crystals and crystals from screening.

|  | Designed Crystals<br>(This work) | Screened Crystals |
| --- | --- | --- |
| Crystal Packing | Sequence encoded | Both sequence and screening<br>condition dependent |
| Crystal Contacts | Distinct, hydrophobic, specific | Continuous, polar, nonspecific |
| Space Group<br>preference | Highly symmetrical (Cubic) | Lower symmetry (monoclinic etc.) |
| Solvent Content | High (80-90%) | 27-65% on average<sup>67</sup> |
| Porosity | Pores within and between cages | Usually uniform pores |
| Crystallization<br>Concentration | Down to 0.2 mg/mL | Usually above 10 mg/mL |
| Crystallization<br>Precipitants | Can crystallize upon mixing<br>without additional precipitants | Usually need high concentration of<br>precipitants |
| Thermal Stability | High (stable up to 95 °C) | Usually unstable above room<br>temperature |

## References

1. Berman, H. M. The Protein Data Bank. Nucleic Acids Res. 28, 235–242 (2000).

2. Hartje, L. F. & Snow, C. D. Protein crystal based materials for nanoscale applications in medicine and biotechnology. WIREs Nanomedicine Nanobiotechnology 11, (2019).

3. Orehek, J., Teslić, D. & Likozar, B. Continuous Crystallization Processes in Pharmaceutical Manufacturing: A Review. Org. Process Res. Dev. 25, 16–42 (2021).

4. Basu, S. K., Govardhan, C. P., Jung, C. W. & Margolin, A. L. Protein crystals for the delivery of biopharmaceuticals. Expert Opin. Biol. Ther. 4, 301–317 (2004).

5. Conejero-Muriel, M., Rodrgíuez-Ruiz, I., Verdugo-Escamilla, C., Llobera, A. & Gavira, J. A. Continuous Sensing Photonic Lab-on-a-Chip Platform Based on Cross-Linked Enzyme Crystals. Anal. Chem. 88, 11919–11923 (2016).

6. Jegan Roy, J. & Emilia Abraham, T. Strategies in Making Cross-Linked Enzyme Crystals. Chem. Rev. 104, 3705–3722 (2004).

7. Heater, B. S., Yang, Z., Lee, M. M. & Chan, M. K. In Vivo Enzyme Entrapment in a Protein Crystal. J. Am. Chem. Soc. 142, 9879–9883 (2020).

8. Mcpherson, A. Introduction to protein crystallization. Methods 34, 254–265 (2004).

9. Desiraju, G. R. Crystal Engineering: A Holistic View. Angew. Chem. Int. Ed. 46, 8342–8356 (2007).

10. Rupp, B. Biomolecular crystallography: principles, practice, and application to structural biology. (Garland Science, 2010).

11. Brodin, J. D. et al. Metal-directed, chemically tunable assembly of one-, two- and three-dimensional crystalline protein arrays. Nat. Chem. 4, 375–382 (2012).

12. Sontz, P. A., Bailey, J. B., Ahn, S. & Tezcan, F. A. A Metal Organic Framework with Spherical Protein Nodes: Rational Chemical Design of 3D Protein Crystals. J. Am. Chem. Soc. 137, 11598– 11601 (2015).

13. Subramanian, R. H. et al. Design of metal-mediated protein assemblies via hydroxamic acid functionalities. Nat. Protoc. 16, 3264–3297 (2021).

14. Sinclair, J. C., Davies, K. M., Vénien-Bryan, C. & Noble, M. E. M. Generation of protein lattices by fusing proteins with matching rotational symmetry. Nat. Nanotechnol. 6, 558–562 (2011).

15. Kostiainen, M. A. et al. Electrostatic assembly of binary nanoparticle superlattices using protein cages. Nat. Nanotechnol. 8, 52–56 (2013).

16. Liljeström, V., Mikkilä, J. & Kostiainen, M. A. Self-assembly and modular functionalization of three-dimensional crystals from oppositely charged proteins. Nat. Commun. 5, 4445 (2014).

17. Brodin, J. D., Auyeung, E. & Mirkin, C. A. DNA-mediated engineering of multicomponent enzyme crystals. Proc. Natl. Acad. Sci. 112, 4564–4569 (2015).

18. Zhou, K. et al. On-Axis Alignment of Protein Nanocage Assemblies from 2D to 3D through the Aromatic Stacking Interactions of Amino Acid Residues. ACS Nano 12, 11323–11332 (2018).

19. Lanci, C. J. et al. Computational design of a protein crystal. Proc. Natl. Acad. Sci. 109, 7304– 7309 (2012).

20. Huang, P.-S., Boyken, S. E. & Baker, D. The coming of age of de novo protein design. Nature 537, 320–327 (2016).

21. Bai, Y., Luo, Q. & Liu, J. Protein self-assembly via supramolecular strategies. Chem. Soc. Rev. 45, 2756–2767 (2016).

22. Luo, Q., Hou, C., Bai, Y., Wang, R. & Liu, J. Protein Assembly: Versatile Approaches to Construct Highly Ordered Nanostructures. Chem. Rev. 116, 13571–13632 (2016).

23. Zhu, J. et al. Protein Assembly by Design. Chem. Rev. 121, 13701–13796 (2021).

24. Leman, J. K. et al. Macromolecular modeling and design in Rosetta: recent methods and frameworks. Nat. Methods 17, 665–680 (2020).

25. Cotton, F. A. Chemical applications of group theory. (Wiley, 1990).

26. Yeates, T. O., Liu, Y. & Laniado, J. The design of symmetric protein nanomaterials comes of age in theory and practice. Curr. Opin. Struct. Biol. 39, 134–143 (2016).

27. Laniado, J. & Yeates, T. O. A complete rule set for designing symmetry combination materials from protein molecules. Proc. Natl. Acad. Sci. 117, 31817–31823 (2020).

28. Fallas, J. A. et al. Computational design of self-assembling cyclic protein homo-oligomers. Nat. Chem. 9, 353–360 (2017).

29. King, N. P. et al. Accurate design of co-assembling multi-component protein nanomaterials. Nature 510, 103–108 (2014).

30. Boyken, S. E. et al. De novo design of protein homo-oligomers with modular hydrogen-bond network–mediated specificity. Science 352, 680–687 (2016).

31. King, N. P. et al. Computational Design of Self-Assembling Protein Nanomaterials with Atomic Level Accuracy. Science 336, 1171–1174 (2012).

32. Sheffler, W. et al. Fast and versatile sequence-independent protein docking for nanomaterials design using RPXDock. http://biorxiv.org/lookup/doi/10.1101/2022.10.25.513641 (2022) doi:10.1101/2022.10.25.513641.

33. Bale, J. B. et al. Structure of a designed tetrahedral protein assembly variant engineered to have improved soluble expression: Designed Protein Tetrahedron. Protein Sci. 24, 1695–1701 (2015).

34. Ueda, G. et al. Tailored design of protein nanoparticle scaffolds for multivalent presentation of viral glycoprotein antigens. eLife 9, e57659 (2020).

35. Boyken, S. E. et al. De novo design of tunable, pH-driven conformational changes. Science 364, 658–664 (2019).

36. Brunette, T. et al. Exploring the repeat protein universe through computational protein design. Nature 528, 580–584 (2015).

37. Wulff, G. On the question of speed of growth and dissolution of crystal surfaces. Z. Kristallogr. 34, 449–530 (1901).

38. Jeliazkov, J. R., Robinson, A. C., García-Moreno E., B., Berger, J. M. & Gray, J. J. Toward the computational design of protein crystals with improved resolution. Acta Crystallogr. Sect. Struct. Biol. 75, 1015–1027 (2019).

39. Lee, S. et al. Shape memory in self-adapting colloidal crystals. Nature 610, 674–679 (2022).

40. Künzl e, M., Eckert, T. & Beck, T. Binary Protein Crystals for the Assembly of Inorganic Nanoparticle Superlattices. J. Am. Chem. Soc. 138, 12731–12734 (2016).

41. Tian, Y. et al. Ordered three-dimensional nanomaterials using DNA-prescribed and valence-controlled material voxels. Nat. Mater. 19, 789–796 (2020).

42. Hsia, Y. et al. Design of multi-scale protein complexes by hierarchical building block fusion. Nat. Commun. 12, 2294 (2021).

43. Fleishman, S. J. et al. RosettaScripts: A Scripting Language Interface to the Rosetta Macromolecular Modeling Suite. PLoS ONE 6, e20161 (2011).

44. Schrödinger, LLC. The PyMOL Molecular Graphics System, Version 1.8. (2015).

45. Studier, F. W. Protein production by auto-induction in high-density shaking cultures. Protein Expr. Purif. 41, 207–234 (2005).

46. Schmitt, J., Hess, H. & Stunnenberg, H. G. Affinity purification of histidine-tagged proteins. Mol. Biol. Rep. 18, 223–230 (1993).

47. Tetter, S. & Hilvert, D. Enzyme Encapsulation by a Ferritin Cage. Angew. Chem. 129, 15129– 15132 (2017).

48. Kabsch, W. XDS. Acta Crystallogr. D Biol. Crystallogr. 66, 125–132 (2010).

49. Minor, W., Cymborowski, M., Otwinowski, Z. & Chruszcz, M. HKL-3000: the integration of data reduction and structure solution – from diffraction images to an initial model in minutes. Acta Crystallogr. D Biol. Crystallogr. 62, 859–866 (2006).

50. Winn, M. D. et al. Overview of the CCP 4 suite and current developments. Acta Crystallogr. D Biol. Crystallogr. 67, 235–242 (2011).

51. McCoy, A. J. et al. Phaser crystallographic software. J. Appl. Crystallogr. 40, 658–674 (2007).

52. Adams, P. D. et al. PHENIX : a comprehensive Python-based system for macromolecular structure solution. Acta Crystallogr. D Biol. Crystallogr. 66, 213–221 (2010).

53. Emsley, P. & Cowtan, K. Coot : model-building tools for molecular graphics. Acta Crystallogr. D Biol. Crystallogr. 60, 2126–2132 (2004).

54. Williams, C. J. et al. MolProbity: More and better reference data for improved all-atom structure validation: PROTEIN SCIENCE.ORG. Protein Sci. 27, 293–315 (2018).

55. Punjani, A., Rubinstein, J. L., Fleet, D. J. & Brubaker, M. A. cryoSPARC: algorithms for rapid unsupervised cryo-EM structure determination. Nat. Methods 14, 290–296 (2017).

56. Carragher, B. et al. Leginon: An Automated System for Acquisition of Images from Vitreous Ice Specimens. J. Struct. Biol. 132, 33–45 (2000).

57. Sun, M. et al. Practical considerations for using K3 cameras in CDS mode for high-resolution and high-throughput single particle cryo-EM. J. Struct. Biol. 213, 107745 (2021).

58. Emsley, P., Lohkamp, B., Scott, W. G. & Cowtan, K. Features and development of Coot. Acta Crystallogr. D Biol. Crystallogr. 66, 486–501 (2010).

59. Wang, R. Y.-R. et al. Automated structure refinement of macromolecular assemblies from cryo-EM maps using Rosetta. eLife 5, e17219 (2016).

60. DiMaio, F., Leaver-Fay, A., Bradley, P., Baker, D. & André, I. Modeling Symmetric Macromolecular Structures in Rosetta3. PLoS ONE 6, e20450 (2011).

61. Barad, B. A. et al. EMRinger: side chain–directed model and map validation for 3D cryo-electron microscopy. Nat. Methods 12, 943–946 (2015).

62. Pettersen, E. F. et al. UCSF Chimera?A visualization system for exploratory research and analysis. J. Comput. Chem. 25, 1605–1612 (2004).

63. Pettersen, E. F. et al. UCSF ChimeraX : Structure visualization for researchers, educators, and developers. Protein Sci. 30, 70–82 (2021).

64. Wang, S. et al. The emergence of valency in colloidal crystals through electron equivalents. Nat. Mater. 21, 580–587 (2022).

65. Senesi, A. J. & Lee, B. Small-angle scattering of particle assemblies. J. Appl. Crystallogr. 48, 1172–1182 (2015).

66. Li, T., Senesi, A. J. & Lee, B. Small Angle X-ray Scattering for Nanoparticle Research. Chem. Rev. 116, 11128–11180 (2016).

67. Matthews, B. W. Solvent content of protein crystals. J. Mol. Biol. 33, 491–497 (1968).

