## Supplementray Information for "Accurate Computational Design of 3D Protein Crystals"

**Supplementary Methods**
**Supplementary Figures S1-S13**
**Supplementary Tables S1-S5**

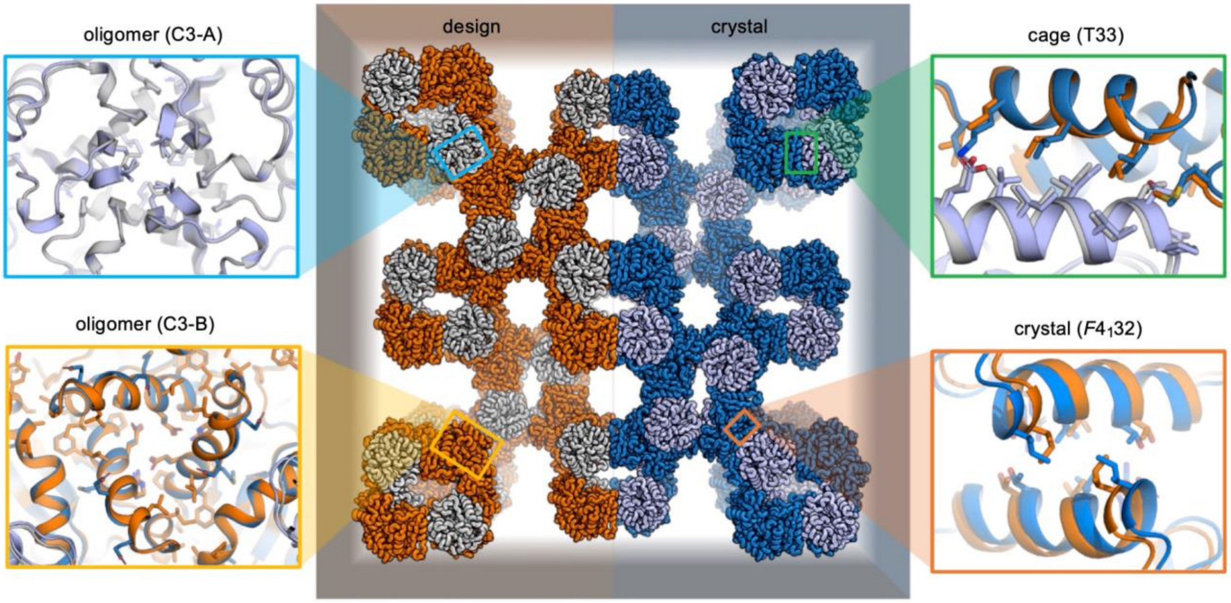


**Supplementary Figure 1. X-ray crystal structure of *F*4$_1$32-1-0 crystal.** Computational design model (rotation corrected (methods), orange and gray) spliced with X-ray crystallography model (skyblue and lightblue) for one unit cell of the crystal. On the corners are close-up views of the four interfaces present in the crystal, design superimposed with crystal model: oligomeric interface of the components on the left, cage interface at right top and crystal interface at right bottom. All-atom RMSD of models: C3-A: 0.69 Å; C3-B: 0.58 Å; T33: 1.06 Å; *F*4$_1$32: 1.81 Å.

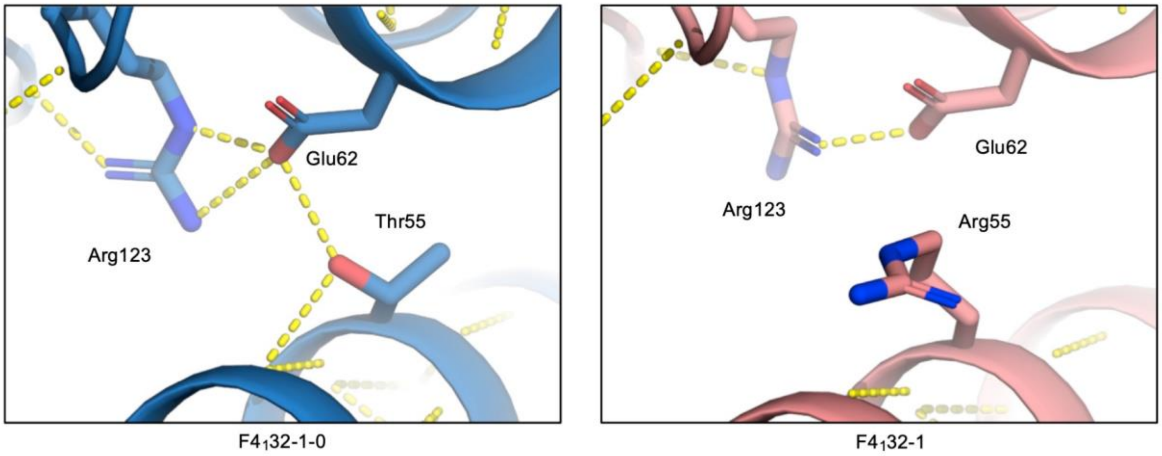


**Supplementary Figure 2. A comparison of crystal structures of *F*4$_1$32-1-0 and *F*4$_1$32-1 around the T55R substitution.** The observed shift in crystal contacts and deviation from the design model is attributed to an undesigned hydrogen bond network around Thr55.

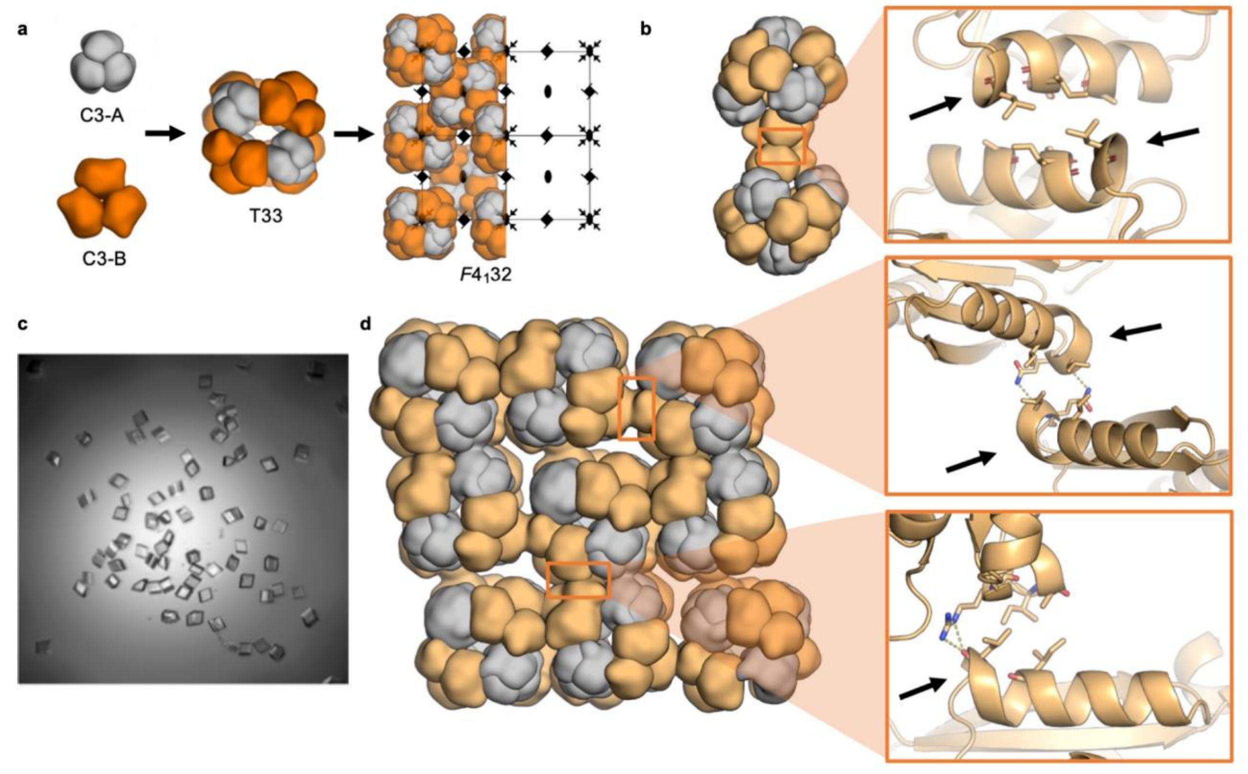


**Supplementary Figure 3. Computational design and experimental characterization of *R*3-1. a,** Overview of the hierarchical design. Symmetry elements of the cage are superimposed with corresponding symmetry elements of the $F4_132$ unit cell. **b,** Design model of adjacent docked T33-15-D3-6 cages and a close-up view of the designed crystal interface. **c,** Optical microscopy images of *R*3-1 crystals formed by mixing purified components at a 1:1 volumetric ratio in 300 mM NaCl, 25 mM Tris pH 8.0, 300 mM imidazole. **d,** Observed R3 crystal packing (light orange and gray) for one unit cell of the crystal. On the corners are close-up views of the two observed crystal contact interfaces present in the crystal. Black arrow points to a helical bulge present at both the designed crystal contact interface and the two observed off-target interfaces.

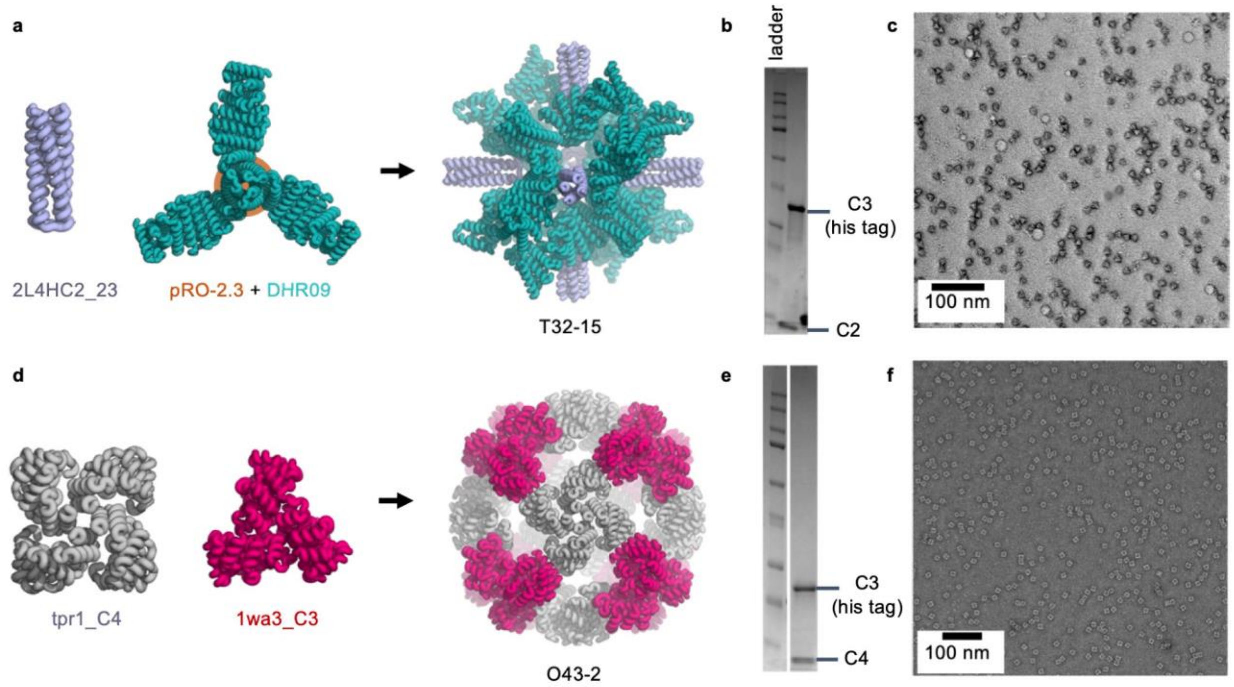


**Supplementary Figure 4. Design scheme and experimental verification of the assembly of new cages. a**, Design model and assembly scheme of T32-15 cage. **b**, SDS-PAGE of IMAC pull-down of protein co-expression of T32-15 cage. **c**, nsEM images of T32-15 cages from IMAC pull-down and SEC purification (assembled *in vivo*). **d**, Design model and assembly scheme of O43-2 cage. **e**, SDS-PAGE of IMAC pull-down of protein co-expression of O43-2 cage. **f**, nsEM images of O43 cages from IMAC pull-down and SEC purification (assembled *in vivo*).

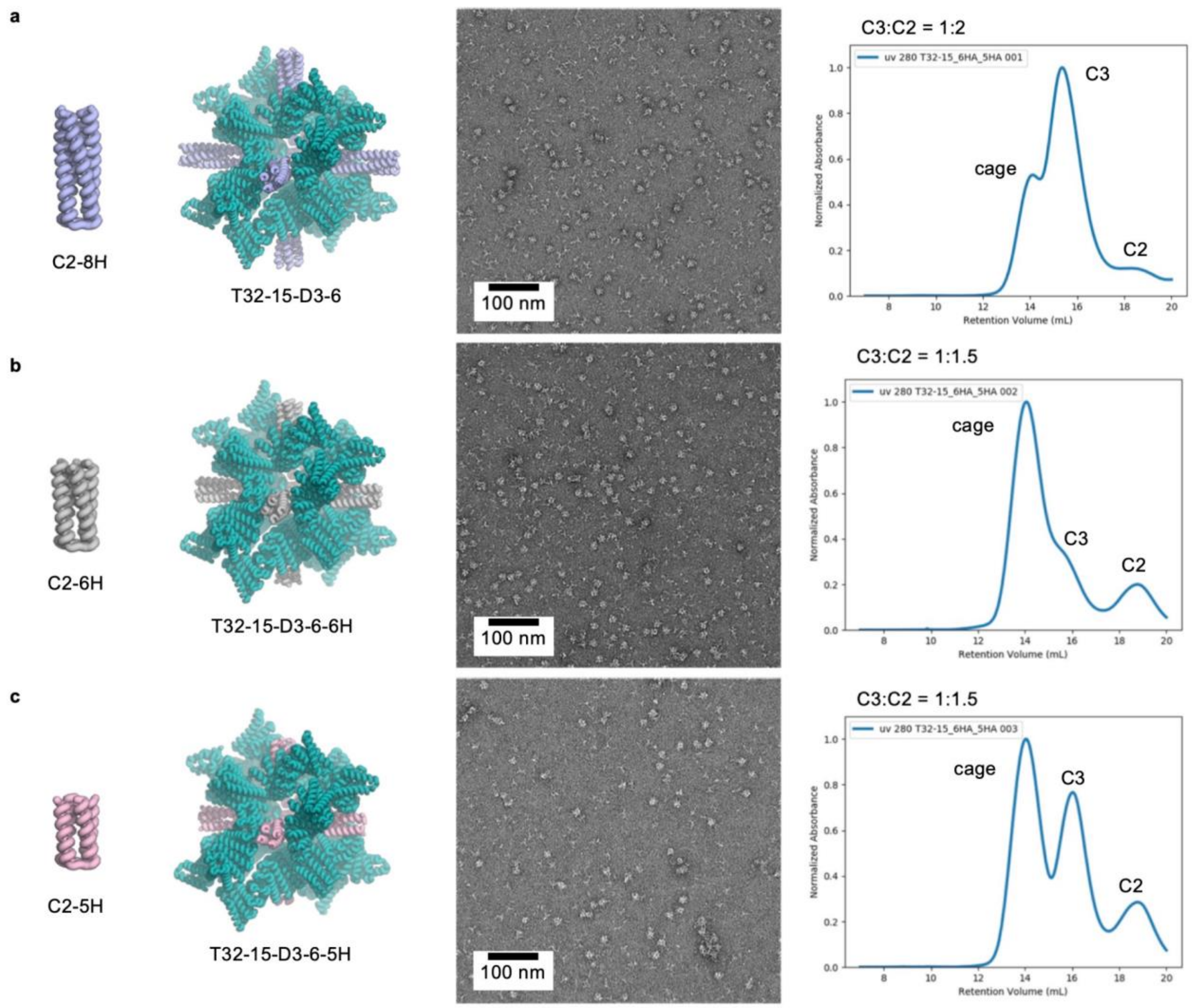


**Supplementary Figure 5. T32-15-D3-6 cage with trimmed C2 component. a,** The original T32-15-D3-6 design with C2 of 8 heptads (C2-8H), nsEM image of equivalent molar mixing and SEC traces. Cage assembly was not complete even with twice the equimolar amount of C2 components. **b,** T32-15-D3-6-6H design with C2 of 6 heptads (C2-6H), nsEM image of equivalent molar mixing and SEC traces. Cage assembly was almost complete with excess C2 components. **c,** T32-15-D3-6-5H design with C2 of 5 heptads (C2-5H), nsEM image of equivalent molar mixing and SEC traces. Incomplete cage assemblies were observed in the presence of excess C2 components.

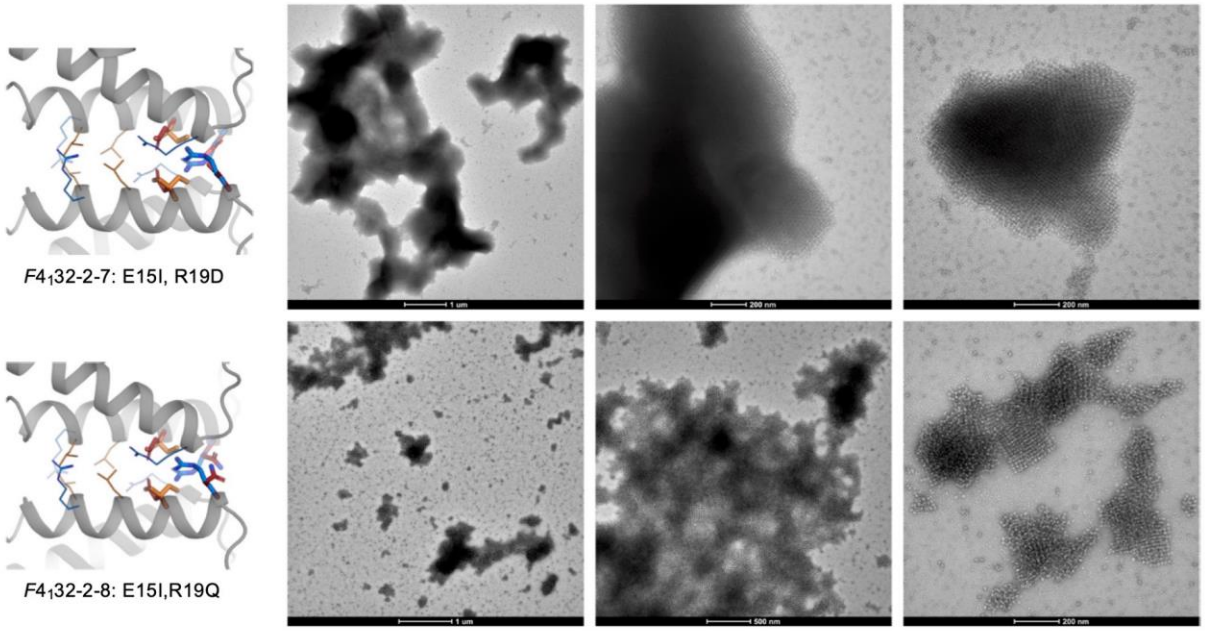


**Supplementary Figure 6. Mutations of *F*4$_1$32-2 crystals into microcrystals and polycrystals**. Models based on crystal structure (left) and their representative crystallization results characterized by nsEM (right).

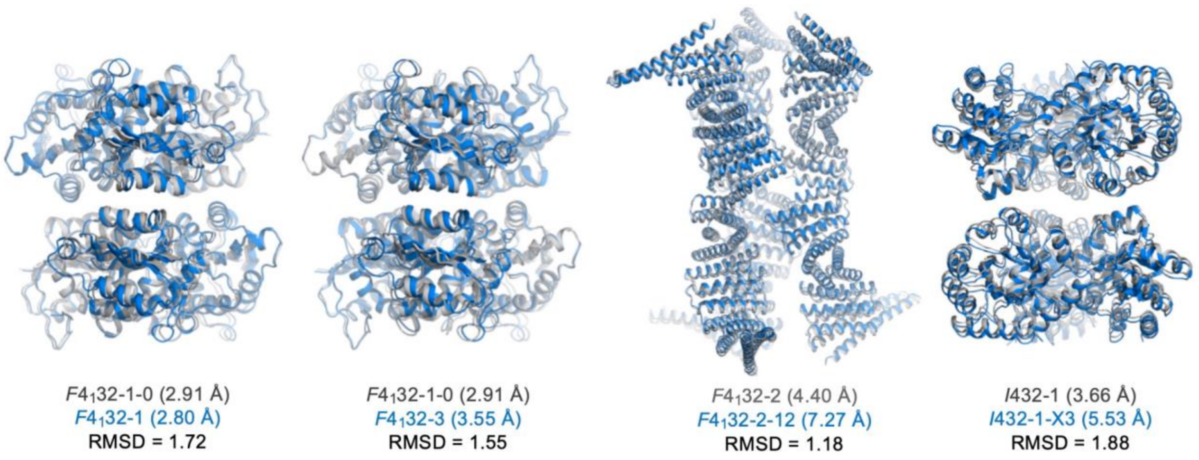


**Supplementary Figure 7. X-ray structures of crystals with mutations as compared to their parent crystal aligned at the dihedral contacts.** Small deviations and shifts of the backbones were observed, potentially due to the flexibility of the cages.

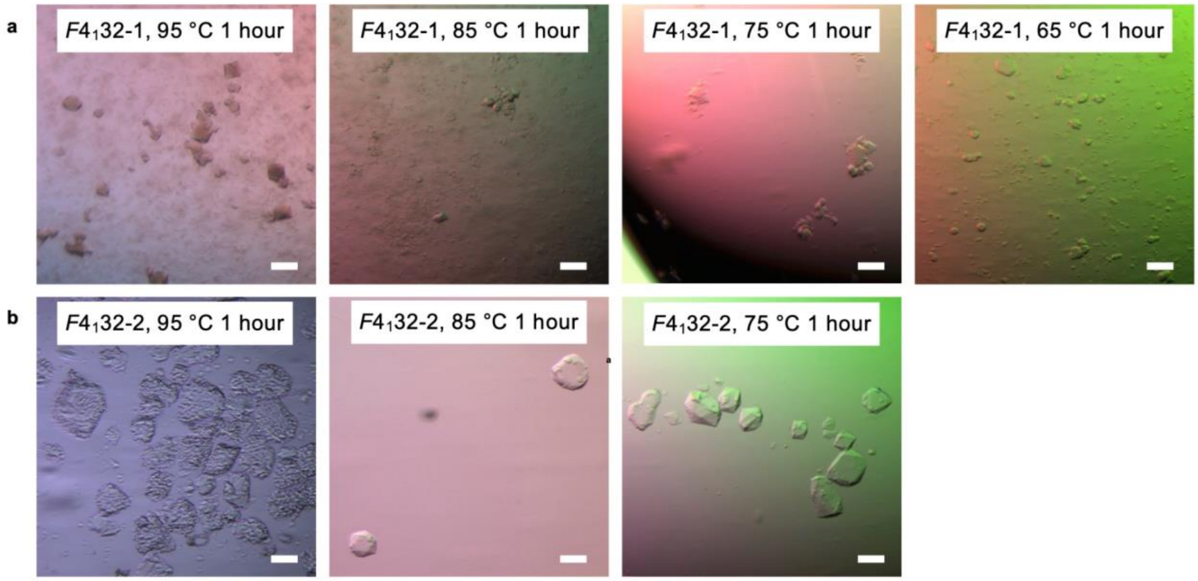


**Supplementary Figure 8. Thermal stability of** $\boldsymbol{F4_{1}32}$**-1 and** $\boldsymbol{F4_{1}32}$**-2 crystals. a**, Optical micrographs of $F4_{1}32$-1 crystals incubated at 95 °C, 85 °C, 75°C, and 65 °C for 1 hour. The crystals were stable up to 65 °C but started to clump together and the colloidal solution became turbid above 75°C. **b**, Optical micrographs of $F4_{1}32$-2 crystals incubated at 95 °C, 85 °C and 75°C for 1 hour. The crystals were stable below 85 °C while dissolved with visible pores at 95°C. See methods for experimental details. Scale bars, 100 µm.

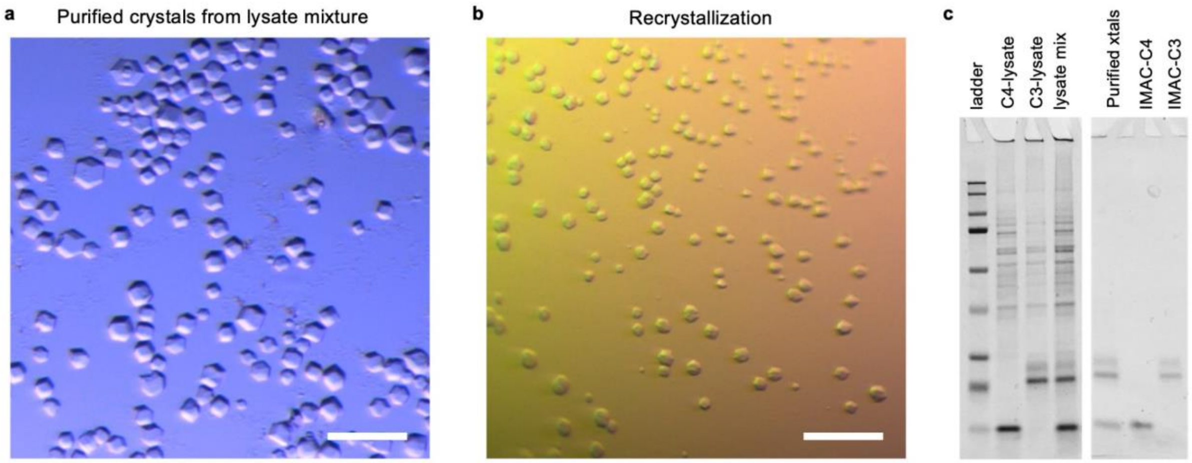


**Supplementary Figure 9. Purification of *I*432-1-CC crystals from lysate mixture. a,** Optical micrograph of crystals purified from mixing lysates (component C4 + C3) overnight and centrifugation. Scale bar, 100 µm. **b**, Purified crystals dissolved in water and recrystallized in 0.5 M NaCl of hanging drop. Scale bar, 100 µm. **c**, SDS-PAGE of proteins from purified crystals. The protein yield is comparable to IMAC purification from the same amount of lysate.

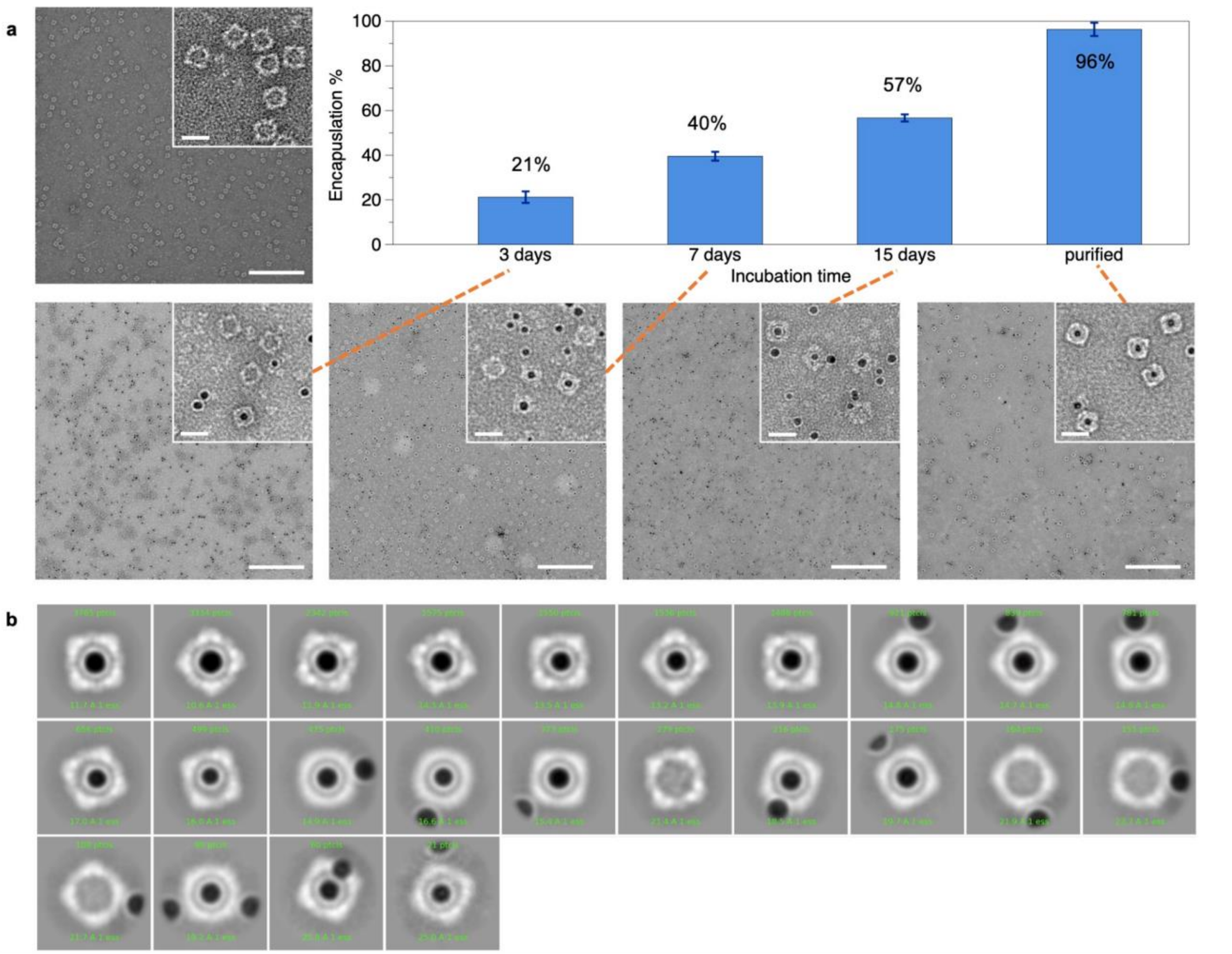


**Supplementary Figure 10. Encapsulation of AuNPs in *I*432-1-CC octahedral cage. a**, The yield of AuNP encapsulation over 15 days of incubation at room temperature. Yield for each sample was estimated by counting the number of empty and encapsulated cages (n = 400-800) in 3-5 nsEM images taken over the grid. Scale bar, 200 nm. Inset, scale bar, 20 nm. **b**, 2D class averages of the nsEM of purified AuNP@*I*432-1-CC cages by CryoSparc.

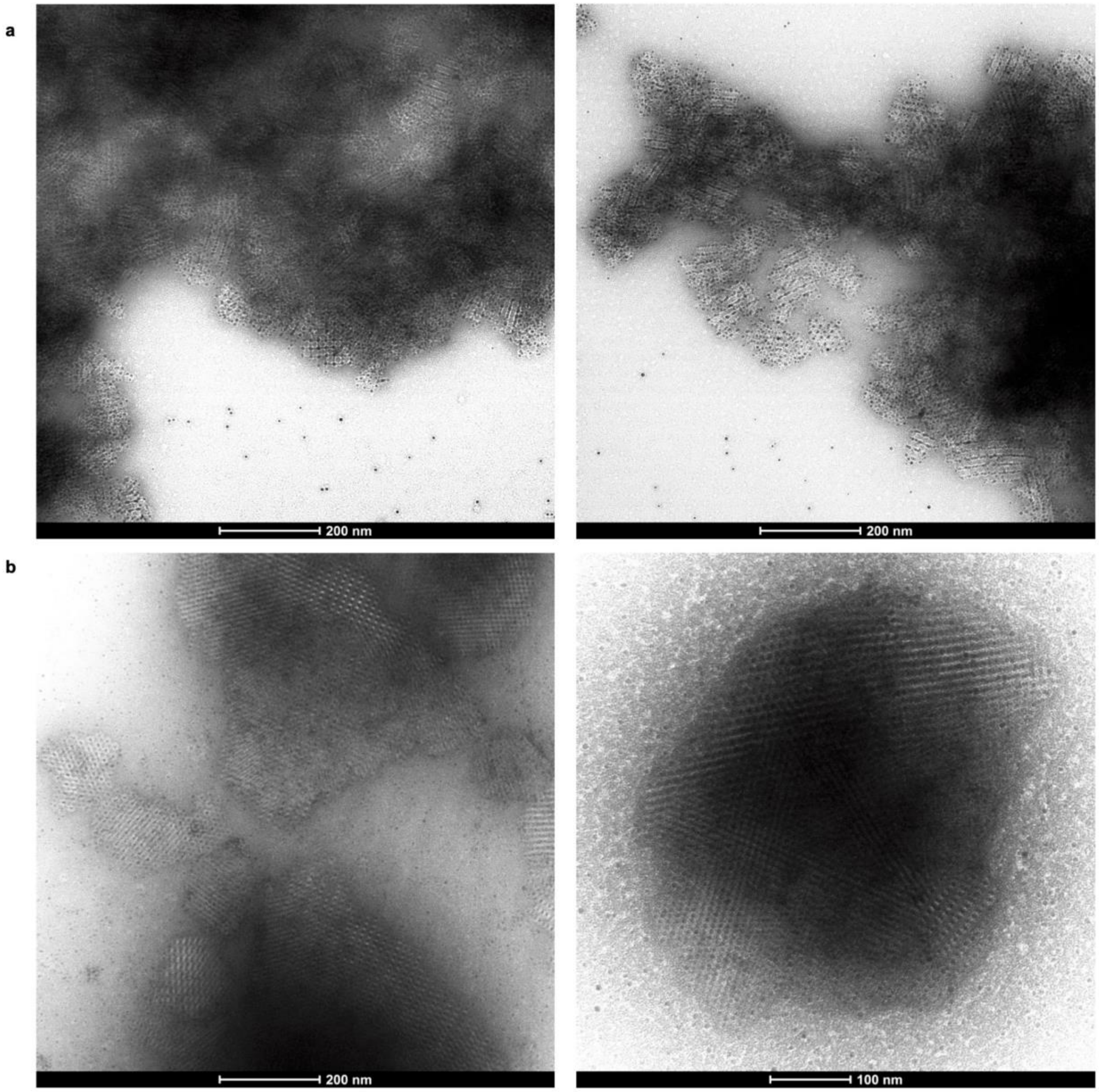


**Supplementary Figure 11. nsEM micrographs of O43-2-D3-6 cage-AuNP coassembly (a) and AuNPs@*I*432-1-CC crystals (b).**

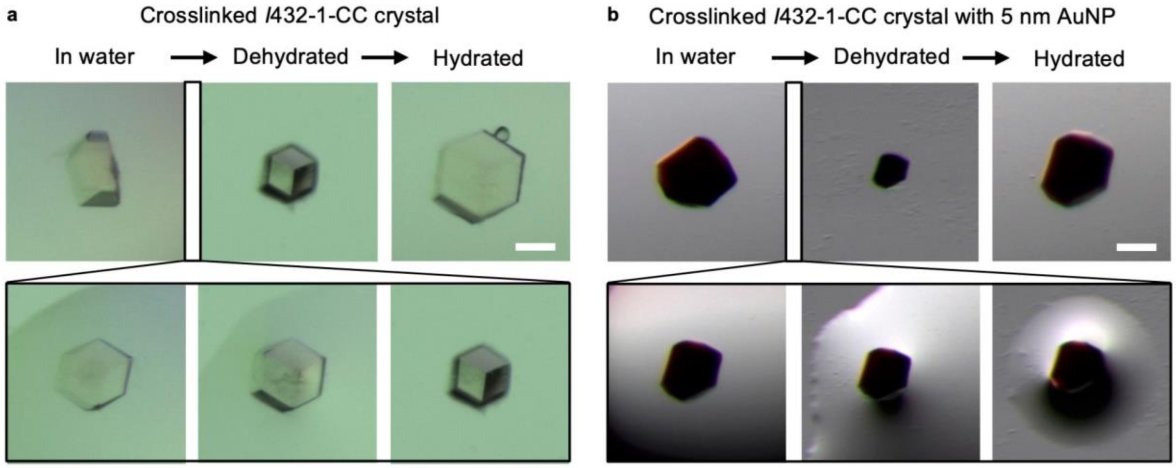


**Supplementary Figure 12. Drying and rehydration cycle of crosslinked crystals, without (a) and with AuNP encapsulation (b).** Both crystals were crosslinked by 2% glutaraldehyde in 0.5 M NaCl overnight. The crystals contracted and expanded evenly while retaining the overall crystal morphology. Scale bars, 50 µm.

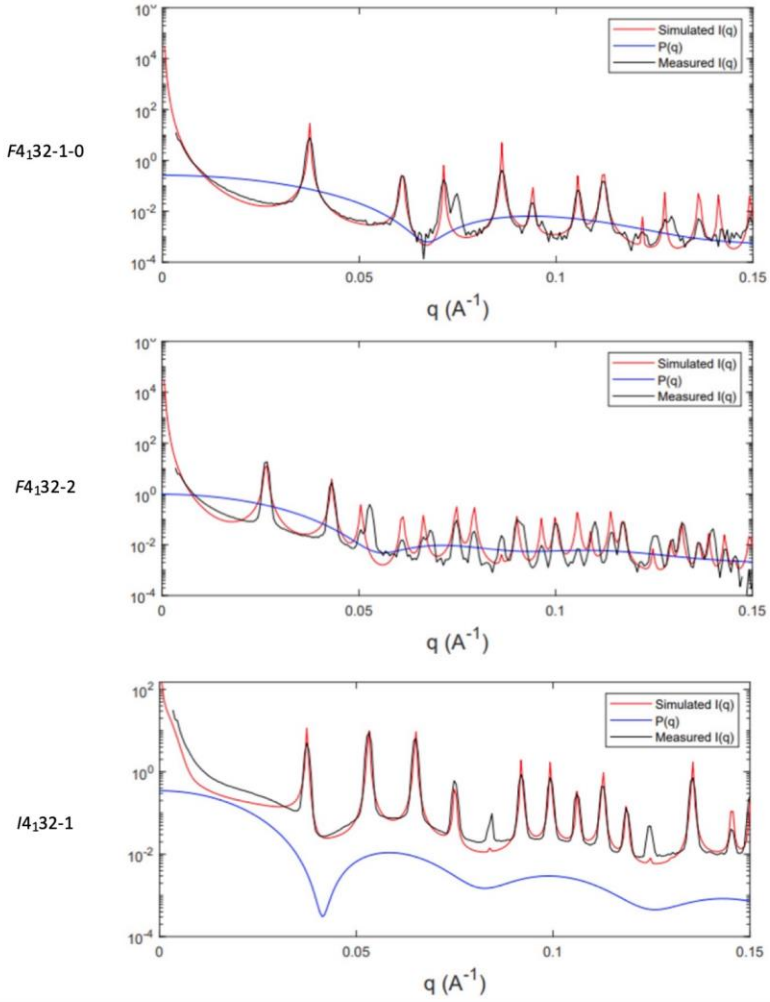


**Supplementary Figure 13. SAXS simulation of protein cage form factor and calculated crystal intensity as compared to experimental data.** P(q) represents the calculated form factor of individual cages.

**Supplementary Table S1.** Crystallization conditions of all designs

"Mixing" refers to directly mixing the two cage components in their purification buffer of 150 mM NaCl, 25 mM Tris pH 8.0. "Hanging drops" and "batch" crystallizations were previously described in the methods section.

| Design name | Method | Cage concentration | NaCl concentration | Time | Crystal Size |
|---|---|---|---|---|---|
| $F4_132$-1-0 | hanging drop | 50 µM | 2.5-3 M | in a week | ~100 µm |
| $F4_132$-1 | mixing | 25 µM | - | overnight | ~100 µm |
| $F4_132$-2-6H | hanging drop | 50 µM | 1-2 M | 3-4 days | ~100 µm |
| $F4_132$-2 | hanging drop | 50 µM | 1-2 M | 3-4 days | ~100 µm |
| $F4_132$-2 | batch | 50-100 µM | 5 M | overnight | ~10 µm |
| $I432$-1 | mixing | 25 µM | - | overnight | ~100 µm |
| $I432$-1 | mixing at 50 mM NaCl | 25 µM | - | in a week | ~100 µm |
| $I432$-1-CC | hanging drop without extra NaCl | 25-100 µM | 0.5 M NaCl | 3-4 days | ~100 µm |
| $I432$-1-CC | batch | 25-50 µM | 0.5-1 M NaCl | minutes | ~10 µm |
| $F4_132$-1-2 | hanging drop | 50 µM | 4 M NaCl | in a week | 20-50 µm |
| $F4_132$-1-3 | hanging drop | 50 µM | 2.5-3 M | in a week | ~100 µm |
| $F4_132$-2-1 | hanging drop | 50 µM | 3 M NaCl | 3-4 days | 20-50 µm |
| $F4_132$-2-3 | hanging drop | 50 µM | 3 M NaCl | 3-4 days | 20-50 µm |
| $F4_132$-2-12 | hanging drop | 50 µM | 0.5 M | 3-4 days | ~100 µm |
| $F4_132$-2-14 | hanging drop | 50 µM | 0.5 NaCl | 3-4 days | 20-50 µm |
| $F4_132$-2-15 | hanging drop | 50 µM | 1 M NaCl | overnight | ~10 µm |
| $I432$-1-7 | mixing | 25 µM | - | minutes | ~10 µm |
| $I432$-1-8 | hanging drop | 25 µM | 1 M NaCl | 3-4 days | 20-50 µm |
| $I432$-1-X3 | hanging drop | 25 µM | 0.5 M NaCl | 3-4 days | ~100 µm |
| $I432$-1-X6 | mixing | 25 µM | - | minutes | ~10 µm |

| | | | | | |
|---|---|---|---|---|---|
| *I*432-1-X9 | hanging drop | 25 µM | 1 M NaCl | 3-4 days | 20-50 µm |
| $F4_1$32-2-ex1 | mixing | 25 µM | - | overnight | 20-50 µm |
| $F4_1$32-2-ex2 | hanging drop | 10-25 µM | 0.5 M NaCl | 3-4 days | 20-50 µm |
| $F4_1$32-2-ex3 | hanging drop | 10-25 µM | 0.5 M NaCl | 3-4 days | 20-50 µm |

**Supplementary Table S2.** Experimental tested mutagenesis of crystal contacts

| Design name | Mutations | Soluble component? | Crystallization? |
|---|---|---|---|
| $F4_132$-1 | T56R | Y | Y, 150 mM NaCl |
| $F4_132$-1-2 | T56A | Y | Y, 4 M NaCl |
| $F4_132$-1-3 | I59V | Y | Y, 3 M NaCl |
| $F4_132$-1-4 | I64V | Y | N, ppts in 5 M NaCl |
| $F4_132$-1-5 | I59V, I64V | Y | N, ppts in 5 M NaCl |
| $F4_132$-1-6 | D63A | N | - |
| $F4_132$-1-7 | T56A, D63M | N | - |
| $F4_132$-1-8 | D63M | N | - |
| $F4_132$-1-9 | K52D | N | - |
| $F4_132$-1-10 | K60E | Y | N, ppts in 5 M NaCl |
| $F4_132$-1-11 | R124D | N | - |
| $F4_132$-1-12 | K50D, R124D | N | - |
| $F4_132$-1-13 | E55K, D63A, D93K | N | - |
| $F4_132$-2-1 | A6S | Y | Y, 3 M NaCl |
| $F4_132$-2-2 | L13V | Y | N, clear |
| $F4_132$-2-3 | L9I | Y | Y, 3 M NaCl |
| $F4_132$-2-4 | L9V | Y | N, clear |
| $F4_132$-2-5 | L9V, L13I | Y | N, clear |
| $F4_132$-2-6 | E16L, R20D | Y | N, ppts |
| $F4_132$-2-7 | E16I, R20D | Y | Y, microcrystals |
| $F4_132$-2-8 | E16I, R20Q | Y | N, polycrystals |
| $F4_132$-2-9 | Ntrim2-4, A6D, E16I, R20Q | Y | N, polycrystals |
| $F4_132$-2-10 | Ntrim2-4, A6D, L9S, E16S, R20Q | Y | N, clear |

| | | | |
|---|---|---|---|
| $F4_132$-2-11 | Ntrim2-4, A6D, L9T, E16I, R20Q | Y | N, clear |
| $F4_132$-2-12 | Ntrim2-4, A6D, E16T, R20Q | Y | Y, 0.5 M NaCl |
| $F4_132$-2-13 | Ntrim2-4, A6D, L9T, E16T, R20Q | Y | N, clear |
| $F4_132$-2-14 | Ntrim2-4, A6D, E16A, R20Q | Y | Y, 0.5 M NaCl |
| $F4_132$-2-15 | Ntrim2-4, A6D, E16T, R20N | Y | Y, 1 M NaCl |
| $F4_132$-2-16 | Ntrim2-4, A6D, E16T, R20D | Y | Y, 3 M NaCl |
| *I*432-2-1 | Q88N, N114N | Y | N, clear |
| *I*432-2-2 | Q88A, N114A | N | - |
| *I*432-2-3 | Q88S, N114S | Y | N, clear |
| *I*432-2-4 | Q88T, N114T | Y | N, clear |
| *I*432-2-5 | Q88L, N114L | Y | N, clear |
| *I*432-2-6 | Q88C, N114C | N | - |
| *I*432-2-7 | Q88K, N114E | Y | Y, 150 mM NaCl |

**Supplementary Table S3.** Protein sequences experimentally tested in this study

*His-tag, Trp for A280 absorbance, and N terminal Met are included in all ordered sequences

| Name | Sequence* | Notes |
|---|---|---|
| T33-15A | MVRGIRGAITVNSDTPTSIIIATILLLEKMLEANGIQSYEELAAV IFTVTEDLTSAFPAEAARQIGMHRVPLLSAREVPVPGSLPRVIRV LALWNTDTPQDRVRHVYLSEAVRLRPDLESAQLEHHHHHH | T33-15 cage C3-A/ $F4_1 32$-1 chain A |
| T33-15B | MSKAKIGIVTVSDRASAGITADISGKAIILALNLYLTSEWEPIYQ VIPDEQDVIETTLIKMADEQDCCLIVTTGGTGPAKRDVTPEATEA VCDRMMPGFGELMRAESLKEVPTAILSRQTAGLRGDSLIVNLPGD PASISDCLLAVFPAIPYCIDLMEGPYLECNEAMIKPFRPKAKLEH HHHHH | T33-15 cage C3-B |
| T33-15-D3-4B | MSKAKIGIVTVSDRASAGITADISGKAIILALNLYLTSEWEPIYQ VIPDEQKVIETTLIKMADIQDCCLIVTTGGTGPAKRDVTPEATEA VCDRMMPGFGELMRAESLKEVPTAILSRQTAGLRGDSLIVNLPGD PASISDCLLAVFPAIPYCIDLMEGPYLECNEAMIKPFRPKAKLEH HHHHH | $F4_1 32$-1-0 chain B |
| T33-15-D3-4B-1 | MSKAKIGIVTVSDRASAGITADISGKAIILALNLYLTSEWEPIYQ VIPDEQKVIERTLIKMADIQDCCLIVTTGGTGPAKRDVTPEATEA VCDRMMPGFGELMRAESLKEVPTAILSRQTAGLRGDSLIVNLPGD PASISDCLLAVFPAIPYCIDLMEGPYLECNEAMIKPFRPKAKLEH HHHHH | $F4_1 32$-1 chain B |
| T33-15-D3-4B-2 | MSKAKIGIVTVSDRASAGITADISGKAIILALNLYLTSEWEPIYQ VIPDEQKVIEATLIKMADIQDCCLIVTTGGTGPAKRDVTPEATEA VCDRMMPGFGELMRAESLKEVPTAILSRQTAGLRGDSLIVNLPGD PASISDCLLAVFPAIPYCIDLMEGPYLECNEAMIKPFRPKAKLEH HHHHH | $F4_1 32$-1-2 chain B |
| T33-15-D3-4B-3 | MSKAKIGIVTVSDRASAGITADISGKAIILALNLYLTSEWEPIYQ VIPDEQKVIETTLVKMADIQDCCLIVTTGGTGPAKRDVTPEATEA VCDRMMPGFGELMRAESLKEVPTAILSRQTAGLRGDSLIVNLPGD PASISDCLLAVFPAIPYCIDLMEGPYLECNEAMIKPFRPKAKLEH HHHHH | $F4_1 32$-1-3 chain B |
| T33-15-D3-4B-10 | MSKAKIGIVTVSDRASAGITADISGKAIILALNLYLTSEWEPIYQ VIPDEQKVIETTLIEMADIQDCCLIVTTGGTGPAKRDVTPEATEA VCDRMMPGFGELMRAESLKEVPTAILSRQTAGLRGDSLIVNLPGD PASISDCLLAVFPAIPYCIDLMEGPYLECNEAMIKPFRPKAKLEH HHHHH | $F4_1 32$-1-10 chain B |

| | | |
|---|---|---|
| T33-15-D3-6B | MSKAKIGIVTVSDRASAGITADISGKAIILALNLYLTSEWEPIYQ<br>VIPDEQDVIEATLSIMADLQDCCLIVTTGGTGPAKRDVTPEATEA<br>VCDRMMPGFGELMRAESLKEVPTAILSRQTAGLRGDSLIVNLPGD<br>PASISDCLLAVFPAIPYCIDLMEGPYLECNEAMIKPFRPKAKLEH<br>HHHHH | *R*3-1 chain B |
| 2L4HC2_23 | TRTEIIRELERSLREQEELAKRLKELLRELERLQREGSSDEDVRE<br>LLREIKELVEEIEKLAREQKYLVEELKRQ | C2 scaffold for T32-15A |
| HFuse_pH192_0046 | DEAEEKARRVAEKVERLKRSGTSEDEIAEEVAREISEVIRTLKES<br>GSSYEVIAEIVARIVAEIVEALKRSGTSEDEIAEIVARVISEVIR<br>TLKESGSSYEVIAEIVARIVAEIVEALKRSGTSEDEIAEIVARVI<br>SEVIRTLKESGSSYEVIAEIVARIVAEIVEALKRSGTSEDEIAEI<br>VARVISEVIRTLKESGSSYEVIAEIVARIVAEIVEALKRSGTSED<br>EIAEIVARVISEVIRTLKESGSSYEVIAEIVARIVAEIVEALKRS<br>GTSEDEIAEIVARVISEVIRTLKESGSSYEVIAEIVARIVAEIVE<br>ALLRSGTSEEEIAKIVARVMNEVLRTLRESGSDFEVIREILRRIL<br>EEINEALKRGGVSEDEIMRIEIKILLMLLRLSTAELERATRSLKA<br>ITEELKKNPSEDALVEHNRAIVEHNRIIVFNNILIALVLEAIVRA<br>IK | C3 scaffold for T32-15B |
| T32-15A | MTRTEIIRELERSLREQEELAKRLMELLLKLLRLQMTGSSDEDVR<br>RLMLRIIELVEEIEELAREQKYLVEELKRQGSWSGLEHHHHHH | T32-15 cage C2 (8 heptads) |
| T32-15B | MGSWSGHHHHHHGSSDEAEEKARRVAEKVERLKRSGTSEDEIAEE<br>VAREISEVIRTLKESGSSYEVIAEIVARIVAEIVEALKRSGTSED<br>EIAEIVARVISEVIRTLKESGSSYEVIAEIVARIVAEIVEALKRS<br>GTSEDEIAEIVARVISEVIRTLKESGSSYEVIAEIVARIVAEIVE<br>ALKRSGTSEDEIAEIVARVISEVIRTLKESGSSAEVIAEIVARIV<br>AEIVEALKRSGTSEDEIAEIVARVISEVIRTLKESGSSSILIALI<br>VARIVAEIVEALKRSGTSEDEIAEIVARVISEVIRTLKESGSSYE<br>IIALIVAMIVAEIVRALLRSGTSEEEIAKIVARVMNEVLRTLRES<br>GSDFEVIREILRLILAAIRAALQKGGVSEDEIMRIEIKILLMLLR<br>LSTAELERATRSLKAITEELKKNPSEDALVEHNRAIVEHNRIIVF<br>NNILIALVLEAIVRAIK | T32-15 cage C3 |
| T32-15-6HA | MRSLREQEELAKRLMELLLKLLRLQMTGSSDEDVRRLMLRIIELV<br>EEIEELAREQKGSWSGLEHHHHHH | T32-15 cage C2 (6 heptads)\<br>$F4_132$-2 chain A |
| T32-15-5HA | MQEELAKRLMELLLKLLRLQMTGSSDEDVRRLMLRIIELVEEIEE<br>GSWSGLEHHHHHH | T32-15 cage C2<br>(5 heptads) |
| T32-15-D3-6B | MDEAEAKALRVALKVEELKRSGTSEDEIAEEVAREISEVIRTLKE<br>SGSSYEVIAEIVARIVAEIVEALKRSGTSEDEIAEIVARVISEVI<br>RTLKESGSSYEVIAEIVARIVAEIVEALKRSGTSEDEIAEIVARV | $F4_132$-2 chain B |

| | | |
|---|---|---|
| | ISEVIRTLKESGSSYEVIAEIVARIVAEIVEALKRSGTSEDEIAE<br>IVARVISEVIRTLKESGSSAEVIAEIVARIVAEIVEALKRSGTSE<br>DEIAEIVARVISEVIRTLKESGSSSILIALIVARIVAEIVEALKR<br>SGTSEDEIAEIVARVISEVIRTLKESGSSYEIIALIVAMIVAEIV<br>RALLRSGTSEEEIAKIVARVMNEVLRTLRESGSDFEVIREILRLI<br>LAAIRAALQKGGVSEDEIMRIEIKILLMLLRLSTAELERATRSLK<br>AITEELKKNPSEDALVEHNRAIVEHNRIIVFNNILIALVLEAIVR<br>AIKGSWGSLEHHHHHH | |
| T32-15-D3-6B-1 | MDEAESKALRVALKVEELKRSGTSEDEIAEEVAREISEVIRTLKE<br>SGSSYEVIAEIVARIVAEIVEALKRSGTSEDEIAEIVARVISEVI<br>RTLKESGSSYEVIAEIVARIVAEIVEALKRSGTSEDEIAEIVARV<br>ISEVIRTLKESGSSYEVIAEIVARIVAEIVEALKRSGTSEDEIAE<br>IVARVISEVIRTLKESGSSAEVIAEIVARIVAEIVEALKRSGTSE<br>DEIAEIVARVISEVIRTLKESGSSSILIALIVARIVAEIVEALKR<br>SGTSEDEIAEIVARVISEVIRTLKESGSSYEIIALIVAMIVAEIV<br>RALLRSGTSEEEIAKIVARVMNEVLRTLRESGSDFEVIREILRLI<br>LAAIRAALQKGGVSEDEIMRIEIKILLMLLRLSTAELERATRSLK<br>AITEELKKNPSEDALVEHNRAIVEHNRIIVFNNILIALVLEAIVR<br>AIKGSWGSLEHHHHHH | $F4_{1}32$-2-1 chain B |
| T32-15-D3-6B-3 | MDEAEAKAIRVALKVEELKRSGTSEDEIAEEVAREISEVIRTLKE<br>SGSSYEVIAEIVARIVAEIVEALKRSGTSEDEIAEIVARVISEVI<br>RTLKESGSSYEVIAEIVARIVAEIVEALKRSGTSEDEIAEIVARV<br>ISEVIRTLKESGSSYEVIAEIVARIVAEIVEALKRSGTSEDEIAE<br>IVARVISEVIRTLKESGSSAEVIAEIVARIVAEIVEALKRSGTSE<br>DEIAEIVARVISEVIRTLKESGSSSILIALIVARIVAEIVEALKR<br>SGTSEDEIAEIVARVISEVIRTLKESGSSYEIIALIVAMIVAEIV<br>RALLRSGTSEEEIAKIVARVMNEVLRTLRESGSDFEVIREILRLI<br>LAAIRAALQKGGVSEDEIMRIEIKILLMLLRLSTAELERATRSLK<br>AITEELKKNPSEDALVEHNRAIVEHNRIIVFNNILIALVLEAIVR<br>AIKGSWGSLEHHHHHH | $F4_{1}32$-2-3 chain B |
| T32-15-D3-6B-7 | MDEAEAKALRVALKVIELKDSGTSEDEIAEEVAREISEVIRTLKE<br>SGSSYEVIAEIVARIVAEIVEALKRSGTSEDEIAEIVARVISEVI<br>RTLKESGSSYEVIAEIVARIVAEIVEALKRSGTSEDEIAEIVARV<br>ISEVIRTLKESGSSYEVIAEIVARIVAEIVEALKRSGTSEDEIAE<br>IVARVISEVIRTLKESGSSAEVIAEIVARIVAEIVEALKRSGTSE<br>DEIAEIVARVISEVIRTLKESGSSSILIALIVARIVAEIVEALKR<br>SGTSEDEIAEIVARVISEVIRTLKESGSSYEIIALIVAMIVAEIV<br>RALLRSGTSEEEIAKIVARVMNEVLRTLRESGSDFEVIREILRLI<br>LAAIRAALQKGGVSEDEIMRIEIKILLMLLRLSTAELERATRSLK<br>AITEELKKNPSEDALVEHNRAIVEHNRIIVFNNILIALVLEAIVR<br>AIKGSWGSLEHHHHHH | $F4_{1}32$-2-7 chain B |
| T32-15-D3-6B-8 | MDEAEAKALRVALKVIELKQSGTSEDEIAEEVAREISEVIRTLKE<br>SGSSYEVIAEIVARIVAEIVEALKRSGTSEDEIAEIVARVISEVI<br>RTLKESGSSYEVIAEIVARIVAEIVEALKRSGTSEDEIAEIVARV<br>ISEVIRTLKESGSSYEVIAEIVARIVAEIVEALKRSGTSEDEIAE | $F4_{1}32$-2-8 chain B |

| | | |
|---|---|---|
| | IVARVISEVIRTLKESGSSAEVIAEIVARIVAEIVEALKRSGTSE<br>DEIAEIVARVISEVIRTLKESGSSSILIALIVARIVAEIVEALKR<br>SGTSEDEIAEIVARVISEVIRTLKESGSSYEIIALIVAMIVAEIV<br>RALLRSGTSEEEIAKIVARVMNEVLRTLRESGSDFEVIREILRLI<br>LAAIRAALQKGGVSEDEIMRIEIKILLMLLRLSTAELERATRSLK<br>AITEELKKNPSEDALVEHNRAIVEHNRIIVFNNILIALVLEAIVR<br>AIKGSWGSLEHHHHHH | |
| T32-15-D3-6B-12 | MEDKALRVALKVTELKQSGTSEDEIAEEVAREISEVIRTLKESGS<br>SYEVIAEIVARIVAEIVEALKRSGTSEDEIAEIVARVISEVIRTL<br>KESGSSYEVIAEIVARIVAEIVEALKRSGTSEDEIAEIVARVISE<br>VIR<br>TLKESGSSYEVIAEIVARIVAEIVEALKRSGTSEDEIAEIVARVI<br>SEVIRTLKESGSSAEVIAEIVARIVAEIVEALKRSGTSEDEIAEI<br>VARVISEVIRTLKESGSSSILIALIVARIVAEIVEALKRSGTSED<br>EIAEIVARVISEVIRTLKESGSSYEIIALIVAMIVAEIVRALLRS<br>GTSEEEIAKIVARVMNEVLRTLRESGSDFEVIREILRLILAAIRA<br>ALQKGGVSEDEIMRIEIKILLMLLRLSTAELERATRSLKAITEEL<br>KKNPSEDALVEHNRAIVEHNRIIVFNNILIALVLEAIVRAIKGSW<br>GSLEHHHHHH | $F4_132$-2-12 chain B |
| T32-15-D3-6B-14 | MEDKALRVALKVAELKQSGTSEDEIAEEVAREISEVIRTLKESGS<br>SYEVIAEIVARIVAEIVEALKRSGTSEDEIAEIVARVISEVIRTL<br>KESGSSYEVIAEIVARIVAEIVEALKRSGTSEDEIAEIVARVISE<br>VIR<br>TLKESGSSYEVIAEIVARIVAEIVEALKRSGTSEDEIAEIVARVI<br>SEVIRTLKESGSSAEVIAEIVARIVAEIVEALKRSGTSEDEIAEI<br>VARVISEVIRTLKESGSSSILIALIVARIVAEIVEALKRSGTSED<br>EIAEIVARVISEVIRTLKESGSSYEIIALIVAMIVAEIVRALLRS<br>GTSEEEIAKIVARVMNEVLRTLRESGSDFEVIREILRLILAAIRA<br>ALQKGGVSEDEIMRIEIKILLMLLRLSTAELERATRSLKAITEEL<br>KKNPSEDALVEHNRAIVEHNRIIVFNNILIALVLEAIVRAIKGSW<br>GSLEHHHHHH | $F4_132$-2-14 chain B |
| T32-15-D3-6B-15 | MEDKALRVALKVTELKNSGTSEDEIAEEVAREISEVIRTLKESGS<br>SYEVIAEIVARIVAEIVEALKRSGTSEDEIAEIVARVISEVIRTL<br>KESGSSYEVIAEIVARIVAEIVEALKRSGTSEDEIAEIVARVISE<br>VIR<br>TLKESGSSYEVIAEIVARIVAEIVEALKRSGTSEDEIAEIVARVI<br>SEVIRTLKESGSSAEVIAEIVARIVAEIVEALKRSGTSEDEIAEI<br>VARVISEVIRTLKESGSSSILIALIVARIVAEIVEALKRSGTSED<br>EIAEIVARVISEVIRTLKESGSSYEIIALIVAMIVAEIVRALLRS<br>GTSEEEIAKIVARVMNEVLRTLRESGSDFEVIREILRLILAAIRA<br>ALQKGGVSEDEIMRIEIKILLMLLRLSTAELERATRSLKAITEEL<br>KKNPSEDALVEHNRAIVEHNRIIVFNNILIALVLEAIVRAIKGSW<br>GSLEHHHHHH | $F4_132$-2-15 chain B |
| T32-15-D3-6B-16 | MEDKALRVALKVTELKDSGTSEDEIAEEVAREISEVIRTLKESGS<br>SYEVIAEIVARIVAEIVEALKRSGTSEDEIAEIVARVISEVIRTL | $F4_132$-2-16 chain B |

| | | |
|---|---|---|
| | KESGSSYEVIAEIVARIVAEIVEALKRSGTSEDEIAEIVARVISEVIRTLKESGSSYEVIAEIVARIVAEIVEALKRSGTSEDEIAEIVARVISEVIRTLKESGSSAEVIAEIVARIVAEIVEALKRSGTSEDEIAEIVARVISEVIRTLKESGSSSILIALIVARIVAEIVEALKRSGTSEDEIAEIVARVISEVIRTLKESGSSYEIIALIVAMIVAEIVRALLRSGTSEEEIAKIVARVMNEVLRTLRESGSDFEVIREILRLILAAIRAALQKGGVSEDEIMRIEIKILLMLLRLSTAELERATRSLKAITEELKKNPSEDALVEHNRAIVEHNRIIVFNNILIALVLEAIVRAIKGSWGSLEHHHHHH | |
| T32-15-D3-6B-ex1 | MEEIVEKAERKLKFLLQEAEEGGKEDALEIAEKLAELAKEALRVLAEAGGSPELMLRLMETAARALARIARLGGDELREEIKKIMAELVAKAISLLIRMLKRSGSSYEEIAEAVAKAVAKIVEAAKESGMSEDEIAEIVARVISEVIRTLKESGSSAEVIAEIVARIVAEIVEALKRSGTSEDEIAEIVARVISEVIRTLKESGSSSILIALIVARIVAEIVEALKRSGTSEDEIAEIVARVISEVIRTLKESGSSYEIIALIVAMIVAEIVRALLRSGTSEEEIAKIVARVMNEVLRTLRESGSDFEVIREILRLILAAIRAALQKGGVSEDEIMRIEIKILLMLLRLSTAELERATRSLKAITEELKKNPSEDALVEHNRAIVEHNRIIVFNNILIALVLEAIVRAIKGSWSGLEHHHHHH | $F4_{1}32$-2-ex1 chain B |
| T32-15-D3-6B-ex2 | MDEKRKRAQKALERAQEALKKGDVEEAVRAAEEAVKAAAEAGDDEMLELVALQAEAIALAAQQQGNDEVKRKAKMVARAAKVAAEIIKLIKELKRNGASYEEIAEEVAKRVAEIVEELKKQGTSEEEIAFIVAAVIAAVIAALKKSGSSAEVIAEIVARIVAEIVEALKRSGTSEDEIAEIVARVISAVIRALKESGSSSILIALIVARIVAEIVEALKRSGTSEDEIAEIVARVISEVIRTLKESGSSYEIIALIVAMIVAEIVRALLRSGTSEEEIAKIVARVMNEVLRTLRESGSDFEVIREILRLILAAIRAALQKGGVSEDEIMRIEIKILLMLLRLSTAELERATRSLKAITEELKKNPSEDALVEHNRAIVEHNRIIVFNNILIALVLEAIVRAIGSWSGLEHHHHHH | $F4_{1}32$-2-ex2 chain B |
| T32-15-D3-6B-ex3 | MDEKRKRAEKALQRAMKALEKGDVEEAVRAAEEAVKAAAEAGDDEMLKKVAAAAELIAMAAQQQGNDEVKRKAKMVARAAKVAAEIIKLIKELKRNGASYEEIAEEVAKRVAEIVEELKKQGTSEEEIAFIVAAVIAAVIAALKKSGSSAEVIAEIVARIVAEIVEALKRSGTSEDEIAEIVARVISAVIRALKESGSSSILIALIVARIVAEIVEALKRSGTSEDEIAEIVARVISEVIRTLKESGSSYEIIALIVAMIVAEIVRALLRSGTSEEEIAKIVARVMNEVLRTLRESGSDFEVIREILRLILAAIRAALQKGGVSEDEIMRIEIKILLMLLRLSTAELERATRSLKAITEELKKNPSEDALVEHNRAIVEHNRIIVFNNILIALVLEAIVRAIKGSWSGLEHHHHHH | $F4_{1}32$-2-ex3 chain B |
| tpr1C4_2 | ALAYVMLGLLLSLLNRLSLAAEAYKKAIELDPNDALAWLLLGSVLEKLKRLDEAAEAYKKAIELKPNDASAWKELGKVLEKLGRLDEAAEAYKKAIELDPEDAEAWKELGKVLEKLGRLDEAAEAYKKAIELDPND | C4 scaffold for O43-2A |

| | | |
|---|---|---|
| 1wa3 | KMEELFKKHKIVAVLRANSVEEAKEKALAVFEGGVHLIEITFTVP DADTVIKELSFLKEKGAIIGAGTVTSVEQCRKAVESGAEFIVSPH LDEEISQFCKEKGVFYMPGVMTPTELVKAMKLGHTILKLFPGEVV GPQFVKAMKGPFPNVKFVPTGGVNLDNVCEWFKAGVLAVGVGSAL VKGTPDEVREKAKAFVEKIRGCTE | C3 scaffold for O43-2B |
| O43-2A | MRGHHHHHHGSSALAYVMLGLLLSLLNRLSLAAEAYKKAIELDPN DALAWLLLGSVLEKLKRLDEAAEAYKKAIELKPNDASAWKELGKV LEKLGRLDEAAKAYAEAIKLDPSDAEAAKELGKVLEKLGQLELAE RAYQLAIELDPND | O43-2 cage C4/ *I*432-1 chain A |
| O43-2B | MRGHHHHHHGSSKMEELFKKHKIVAVLRANSVEEAKEKALAVFRG GVHLIEITFTVPDADTVIKELSFLKEKGAIIGAGTVTSVEQCRKA VESGAEFIVSPHLDEEISQFCKEKGVFYMPGVMTPTELVKAMKLG HTILKLFPGEVVGPQFVKAMKGPFPNVKFVPTGGVNDQNVCEWFK AGVLAVGVGSALVKGTPEQVEMLAVLFVAKIAGCTE | O43-2 cage C3 |
| O43-2-D3-6B | MRGHHHHHHGSSKMEELFKKHKIVAVLRANSVEEAKEKALAVFRG GVHLIEITFTVPDADTVIKELSFLKEKGAIIGAGTVTSLEQCQKA VESGAEFIVSPHLDPEISKFCKINGVFYMPGVMTPTELVKAMKLG HTILKLFPGEVVGPQFVKAMKGPFPNVKFVPTGGVNDQNVCEWFK AGVLAVGVGSALVKGTPEQVEMLAVLFVAKIAGCTE | *I*432-1 chain B |
| O43-2A-CH | MALAYVMLGLLLSLLNRLSLAAEAYKKAIELDPNDALAWLLLGSV LEKLKRLDEAAEAYKKAIELKPNDASAWKELGKVLEKLGRLDEAA KAYAEAIKLDPSDAEAAKELGKVLEKLGQLELAERAYQLAIELDP NDLEHHHHHH | *I*432-1-CC chain A |
| O43-2-D3-6B-CH | MKMEELFKKHKIVAVLRANSVEEAKEKALAVFRGGVHLIEITFTV PDADTVIKELSFLKEKGAIIGAGTVTSLEQCQKAVESGAEFIVSP HLDPEISKFCKINGVFYMPGVMTPTELVKAMKLGHTILKLFPGEV VGPQFVKAMKGPFPNVKFVPTGGVNDQNVCEWFKAGVLAVGVGSA LVKGTPEQVEMLAVLFVAKIAGCTELEHHHHHH | *I*432-1-CC chain B |
| O43-2-D3-6B-7 | MRGHHHHHHGSSKMEELFKKHKIVAVLRANSVEEAKEKALAVFRG GVHLIEITFTVPDADTVIKELSFLKEKGAIIGAGTVTSLEQCKKA VESGAEFIVSPHLDPEISKFCKIEGVFYMPGVMTPTELVKAMKLG HTILKLFPGEVVGPQFVKAMKGPFPNVKFVPTGGVNDQNVCEWFK AGVLAVGVGSALVKGTPEQVEMLAVLFVAKIAGCTE | *I*432-1-7 chain B |
| O43-2-D3-6B-8 | MRGHHHHHHGSSKMEELFKKHKIVAVLRANSVEEAKEKALAVFRG GVHLIEITFTVPDADTVIKELSFLKEKGAIIGAGTVTSLEACRKA VESGAEFIVSPHLDPEISKFCKIAGVFYMPGVMTPTELVKAMKLG HTILKLFPGEVVGPQFVKAMKGPFPNVKFVPTGGVNDQNVCEWFK AGVLAVGVGSALVKGTPEQVEMLAVLFVAKIAGCTE | *I*432-1-13 chain B |
| O43-2-D3-6B-X3 | MRGHHHHHHGSSKMEELFKKHKIVAVLRANSVEEAKEKALAVFRG GVHLIEITFTVPDADTVIKELSFLKEKGAIIGAGTVTDAKECARA VRSGAEFIVSPHLDPEISKFCKAQGVFYMPGVMTPTELVKAMKLG HTILKLFPGEVVGPQFVKAMKGPFPNVKFVPTGGVNDQNVCEWFK | *I*432-1-X3 chain B |

| | | |
|---|---|---|
| | AGVLAVGVGSALVKGTPEQVEMLAVLFVAKIAGCTE | |
| O43-2-D3-6B-X6 | MRGHHHHHHGSSKMEELFKKHKIVAVLRANSVEEAKEKALAVFRG GVHLIEITFTVPDADTVIKELSFLKEKGAIIGAGTVTDKRQCKKA VESGAEFIVSPHLDPEISEFCKMEGVFYMPGVMTPTELVKAMKLG HTILKLFPGEVVGPQFVKAMKGPFPNVKFVPTGGVNDQNVCEWFK AGVLAVGVGSALVKGTPEQVEMLAVLFVAKIAGCTE | *I*432-1-X6 chain B |
| O43-2-D3-6B-X9 | MRGHHHHHHGSSKMEELFKKHKIVAVLRANSVEEAKEKALAVFRG GVHLIEITFTVPDADTVIKELSFLKEKGAIIGAGTVTDKRQCKKA VESGAEFIVSPHLDPEISEFCKMMGVFYMPGVMTPTELVKAMKLG HTILKLFPGEVVGPQFVKAMKGPFPNVKFVPTGGVNDQNVCEWFK AGVLAVGVGSALVKGTPEQVEMLAVLFVAKIAGCTE | *I*432-1-X9 chain B |

**Supplementary Table S4.** Crystal structure data collection and refinement statistics

| | **$F4_132$-1-0 (8CUU)** | **$F4_132$-1 (8CUV)** | **$F4_132$-1-3 (8CUW)** | **R3-1 (8CUX)** |
|---|---|---|---|---|
| **Data Collection** | | | | |
| Space group | F 41 3 2 | F 41 3 2 | F 41 3 2 | R 3 :H |
| Cell parameters | | | | |
| a,b,c (Å) | 289.61, 289.61, 289.61 | 293.55, 293.55, 293.55 | 285.99, 285.99, 285.99 | 50.22, 50.22, 116.00 |
| α, β, γ (°) | 90, 90, 90 | 90, 90, 90 | 90, 90, 90 | 90, 90, 120 |
| Resolution (Å) | 87.32 - 2.91 (3.01 - 2.91) | 49.62 - 2.8 (2.9 - 2.8) | 101.10 - 3.55 (3.67 - 3.55) | 34.8 - 1.85 (1.91 - 1.85) |
| $R_{merge}$ | 0.241 (6.326) | 0.122 (3.125) | 0.258 (3.625) | 0.114 (1.222) |
| $R_{pim}$ | 0.027 (0.700) | 0.032 (0.521) | 0.042 (0.572) | 0.057 (0.618) |
| $I/\sigma(I)$ | 27.00 (0.93) | 22.00 (0.98) | 15.93 (1.06) | 7.73 (1.31) |
| $CC_{1/2}$ | 1.000 (0.403) | 0.999 (0.645) | 0.999 (0.643) | 0.995 (0.663) |
| Completeness (%) | 99.94 (99.52) | 99.17 (94.52) | 99.95 (99.68) | 99.94 (100.00) |
| Redundancy | 79.4 (82.9) | 39.7 (40.7) | 39.2 (41.5) | 4.9 (4.9) |
| **Refinement** | | | | |
| Resolution (Å) | 87.32 - 2.91 (3.01 - 2.91) | 49.62 - 2.8 (2.9 - 2.8) | 101.10 - 3.55 (3.67 - 3.55) | 34.8 - 1.85 (1.91 - 1.85) |
| No. reflections | 23382 (2286) | 26999 (2537) | 12632 (1235) | 9321 (935) |
| $R_{work}$ / $R_{free}$ (%) | 19.28 (31.57)/ 20.40 (36.33) | 19.57 (28.63)/ 22.35 (34.31) | 22.40 (33.47)/ 24.80 (39.34) | 21.44 (27.40) /26.21 (31.97) |
| No. atoms | | | | |
| Protein | 2252 | 2207 | 2232 | 932 |
| Ramachandran Favored/allowed/ Outlier (%) | 95.21/4.45/0.34 | 97.55/2.45 /0.00 | 92.76/6.90 /0.34 | 96.58/3.42 /0.00 |
| R.m.s. deviations | | | | |
| Bond lengths (Å) | 0.008 | 0.002 | 0.004 | 0.007 |
| Bond angles (˚) | 1.08 | 0.50 | 0.66 | 0.84 |
| $B_{factors}$ (Å$^2$) | | | | |
| Protein | 107.63 | 39.12 | 156.87 | 26.07 |

| | **$F4_132$-2 (8CWS)** | **$F4_132$-2-12** | **I432-1(NaCl)(8CUS)** | **I432-1(Imd) (8CUT)** | **I432-1-X3 (8CWZ)** |
|---|---|---|---|---|---|
| **Data Collection** | | | | | |
| Space group | F 41 3 2 | F 41 3 2 | I 4 3 2 | I 4 3 2 | I 4 3 2 |
| Cell parameters | | | | | |
| a,b,c (Å) | 405.41, 405.41, 405.41 | 403.65, 403.65, 403.65 | 235.72, 235.72, 235.72 | 236.47, 236.47, 236.47 | 239.29, 239.29, 239.29 |
| α, β, γ (°) | 90, 90, 90 | 90, 90, 90 | 90, 90, 90 | 90, 90, 90 | 90, 90, 90 |
| Resolution (Å) | 47.78 - 4.4 (4.55 - 4.4) | 121.70 - 7.37 (7.63 - 7.37) | 48.12 - 3.98 (4.12 - 3.98) | 46.38 - 4.00 (4.14 - 4.00) | 169.2 - 5.53 (5.72 - 5.53) |
| $R_{merge}$ | 0.381 | 0.1701 (1.716) | 0.193 (3.678) | 0.054 (0.165) | 0.104 (1.169) |
| $R_{pim}$ | 0.381 | 0.040 (0.366) | 0.031 (0.577) | 0.05457 (0.1658) | 0.023 (0.547) |
| *I/σ(I)* | 1.00 | 18.73 (2.60) | 24.51 (1.95) | 22.06 (2.95) | 23.35 (3.96) |
| $CC_{1/2}$ | 0.995 | 0.796 (0.847) | 1.000 (0.680) | 0.991 (0.917) | 0.997 (0.861) |
| Completeness (%) | 99.46 (100.00) | 99.93 (100.00) | 99.83 (100.00) | 99.41 (96.46) | 99.2 (97.3) |
| Redundancy | 2.0 (2.0) | 20.9 (22.8) | 39.5 (41.1) | 1.8 (1.3) | 20.9 (15.4) |
| **Refinement** | | | | | |
| Resolution (Å) | 47.78 - 4.4 (4.55 - 4.4) | 121.70 - 7.37 (7.63 - 7.37) | 48.12 - 3.98 (4.12 - 3.98) | 46.38 - 4.00 (4.14 - 4.00) | 169.2 - 5.53 (5.72 - 5.53) |
| No. reflections | 18577 (1819) | 4158 (403) | 9910 (969) | 9814 (926) | 3415 (243) |
| $R_{work}$ / $R_{free}$ (%) | 33.59 (41.67)/ 35.14 (40.63) | 23.29 (28.79)/ 27.88 (28.15) | 17.71 (21.48)/ 21.66 (29.71) | 18.75 (26.23)/ 22.49 (30.94) | 24.11 (36.46)/ 28.72 (29.03) |
| No. atoms | | | | | |
| Protein | 3930 | 3906 | 2552 | 2560 | 2562 |
| Ramachandran Favored/allowed/ Outlier (%) | 99.20/0.60/0.20 | 98.40/1.60 /0.00 | 93.07/6.02/0.90 | 96.70/3.30/0.00 | 87.43/9.88 /2.69 |
| R.m.s. deviations | | | | | |
| Bond lengths (Å) | 0.002 | 0.003 | 0.003 | 0.002 | 0.008 |
| Bond angles (˚) | 0.37 | 0.55 | 0.65 | 0.40 | 1.21 |
| $B_{factors}$ (Å$^2$) | | | | | |
| Protein | 170.49 | 484.38 | 61.06 | 71.57 | 286.140 |

**Supplementary Table S5.** T32-15 CryoEM Data Collection Statistics

| | **T32-15** |
|---|---|
| Microscope | Titan Krios |
| Voltage (kV) | 300 |
| Detector | Gatan K3 Summit |
| Recording mode | Super Resolution |
| Magnification | 22,500 X |
| Movie micrograph pixel size (Å) | 0.5144 |
| Dose rate ($e^{-}$/Å$^2$/s) | 20 |
| No. of frames per movie micrograph | 50 |
| Frame exposure time (ms) | 50 |
| Movie micrograph exposure time (s) | 2.5 |
| Total dose ($e^{-}$/Å$^2$) | 50 |
| Under focus range (μm) | 0.8 - 2.2 |
| Number of movie micrographs | 4202 |
